# ScRNA-seq and scATAC-seq reveal that sertoli cell mediate spermatogenesis disorders through stage-specific communications in non-obstructive azoospermia

**DOI:** 10.1101/2024.04.09.588807

**Authors:** Shimin Wang, Hongxian Wang, Bicheng Jin, Hongli Yan, Qingliang Zheng, Dong Zhao

**Affiliations:** Prenatal Diagnosis Center, The Eighth Affiliated Hospital, Sun Yat-sen University, 3025# Shennan Road, Shenzhen 518000, China; Department of Gynaecology and Obstetrics, Ninth People’s Hospital, Shanghai Jiao Tong University School of Medicine, 639 Zhizaoju Road, Shanghai 200011, China; Department of Urology and Andrology, School of Medicine, Renji Hospital, Shanghai Jiao Tong University, 160 Pujian Road, Pudong New Area, Shanghai, 200127, China; Department of Surgical Subject, Guizhou Electric Staff Hospital, No. 22 Dongtan Lane, Guanshui Road, Nanming District, Guiyang City, Guizhou Province, 550002, China; Reproductive medicine center, the Navy Medical University, 168 Changhai Road, Yangpu District, Shanghai, 200082, China

**Keywords:** Non-obstructive azoospermia, Single cell RNA-sequencing, Single cell ATAC-sequencing, Germ cell, Sertoli cell

## Abstract

Non-obstructive azoospermia (NOA) belongs to male infertility due to spermatogenesis failure. However, evidence for cell type-specific abnormalities of spermatogenesis disorders in NOA remains lacking. We performed single-cell RNA sequencing (scRNA-seq) and single-cell assay for transposase-accessible chromatin sequencing (scATAC-seq) on testicular tissues from patients with obstructive azoospermia(OA) and NOA. HE staining confirmed the structural abnormalities of the seminiferous tubules in NOA patients. We identified 12 germ cell subtypes (spermatogonial stem cell-0 (SSC0), SSC1, SSC2, diffing-spermatogonia (Diffing-SPG), diffed-spermatogonia (Diffed-SPG), pre-leptotene (Pre-Lep), leptotene-zygotene (L-Z), pachytene (Pa), diplotene (Di), spermatids-1 (SPT1), SPT2, and SPT3) and 8 Sertoli cell subtypes (SC1-SC8). Among them, three novel Sertoli cell subtypes phenotypes were identified, namely SC4/immature, SC7/mature, and SC8/further mature Sertoli cells. For each germ or Sertoli cell subtype, we identified unique new markers, among which immunofluorescence confirmed co-localization of ST3GAL4, A2M, ASB9, and TEX19 and DDX4 (classical marker of germ cell). PRAP1, BST2, and CCDC62 were co-expressed with SOX9 (classical marker of Sertoli cell) in testes tissues also confirmed by immunofluorescence. The interaction between germ cell subtypes and Sertoli cell subtypes exhibits stage-specific-matching pattern, as evidenced by SC1/2/5/7 involving in SSC0-2 development, SC3 participating in the whole process of spermiogenesis, SC4/6 participating in Diffing and Diffed-SPG development, and SC8 involving in the final stage of SPT3. This pattern of specific interactions between subtypes of germ cell and Sertoli cell was confirmed by immunofluorescence of novel markers in testes tissues. The interaction was mainly contributed by Notch1/2/3 signaling. Our study profiled the single-cell transcriptome of human spermatogenesis and provided many potentials molecular markers for developing testicular puncture specific marker kits for NOA patients.

**Impact statement:** The specific interactions between Sertoli cell subtypes and different stages of germ subgroups maintain normal development of germ cells. SC3 subtype absence may be the key factor leading to NOA.

## Introduction

Infertility is a common disease affecting approximately 15% of couples, and 50% of cases in infertility is attributed to male factor infertility (Agarwal, Mulgund, Hamada, & Chyatte, 2015). Male infertility is mainly associated with sexual dysfunction, endocrine, varicocele, and reproductive system infection(Naz & Kamal, 2017).Azoospermia includes obstructive azoospermia (OA) and non-obstructive azoospermia (NOA). NOA accounts for about 1% of adult males and 10%-15% of infertile males (Fakhro et al., 2018). The etiology of NOA is varied, including cryptorchidism, post-pubertal mumps orchitis, and prior testicular torsion (Yao et al., 2022). At present, assisted reproductive technology, such as microdissection testicular sperm extraction and intracytoplasmic sperm injection, has made certain progress, providing help for patients to obtain biological offspring. At the same time, spermatogonial stem cell culture and transplantation technology and testicular tissue transplantation technology has also made some progress, but the clinical application is still not mature (Gassei & Orwig, 2016). At present, the etiology and mechanism of many NOA patients are still unknown, which poses a challenge to treatment. Therefore, it is particularly urgent to further study the molecular mechanisms affecting spermatogenesis in NOA patients.

The formation of mature spermatozoa can be delineated into three stages: the proliferation of spermatogonial cells, meiotic division of spermatocytes, and the maturation of sperm cells (Jegou, 1993). The seminiferous tubule comprises three distinct cell types: Sertoli cells, extending from the basal membrane to the tubular lumen; Germ cells of various generations; Peritubular cells, surrounding sertoli cells and germ cells, isolated from sertoli cells by the extracellular matrix. In most mammals, germ cells undergo continuous renewal, while sertoli cells cease division during the pubertal developmental period, forming the seminiferous epithelium. The lining of the seminiferous epithelium is uniquely intricate within a complex tissue structure. At any given point in a seminiferous tubule, several generations of germ cells develop simultaneously in the process of contacting sertoli cells from the basal epithelium to the apex. The evolution of each generation of germ cells is strictly synchronized with others, resulting in the formation of specific cell associations or stages at a particular segment of the tubule. The complete temporal sequence of these cell associations or stages until the reappearance of the initial association in a defined region of the tubule is referred to as the seminiferous epithelial cycle (Hess & Moore, 1993). Adult spermatogonial stem cells are distributed along the basal membrane at the base of the seminiferous tubule, necessitating a delicate balance between self-renewal and differentiation. Throughout this process, only sertoli cells consistently maintain physical contact with sperm cells. Consequently, we posit that the entire process of spermatogenesis inevitably involves intense information exchange with the occurrence of sertoli cells. However, the specifics of this interactive process remain incompletely understood at present.

Single-cell transcriptome sequencing (scRNA-seq) technology is used for high-throughput molecular detection of a single cell and exploring the whole gene expression profiles of a single cell (Tan, Song, & Wilkinson, 2020; Zhao et al., 2020). It can delineate the existence of rare cell subsets and the degree of cellular heterogeneity (Tirumalasetty, Bhattacharya, Mohiuddin, Baki, & Choubey, 2024; Vertesy et al., 2018). Liao et al. (2019) performed scRNA-seq analysis on mouse testicular germ cells at 5.5 days after birth, revealing the heterogeneity of gene expression in undifferentiated spermatogonial cells. Guo et al. (2018) performed scRNA-seq of testicular cells from young adults and found that there were five different spermatogonial states accompanying human spermatogonial differentiation. In addition, scATAC-seq is a widely used method to obtain a genome-wide snapshot of chromatin accessibility, signatures of active transcription and transcription factor (TF) binding (Mimitou et al., 2021). Cellular identity is strongly affected by the epigenetic wiring of the cell, which is measured by scATAC-seq (Pervolarakis et al., 2020). Therefore, scRNA-seq and scATAC-seq can be used to analyze spermatogenesis.

In this study, we performed an integrative analysis of scRNA-seq and scATAC-seq in testicular tissues from two OA and three NOA patients. Cellular heterogeneity and marker gene identification results were analyzed, and the potential functional mechanism was inferred. Altogether, our results provide comprehensive understanding of the maturation of the spermatogenic microenvironment and the mechanisms underlying pathogenesis, offering novel targets for NOA treatment strategies.

## Material and methods

### Human sample acquisition

Testicular tissues were obtained from two OA patients (OA1-P1 and OA2-P2) and three NOA patients (NOA1-P3, NOA2-P4, NOA3-P5) using micro-dissection of testicular sperm extraction separately. Notably, the sperm concentration of the three NOA patients varied in descending order (NOA1 > NOA2 > NOA3,Table1), with NOA3 being the patient with complete absence of sperm. Patient clinical and laboratory characteristics are presented in Table 1. This study was approved by the Ethics Committee of School of Medicine, Renji Hospital, Shanghai Jiao Tong University [approval number: KY2020-193]. All patients provided written informed consent.

**Table 1.**
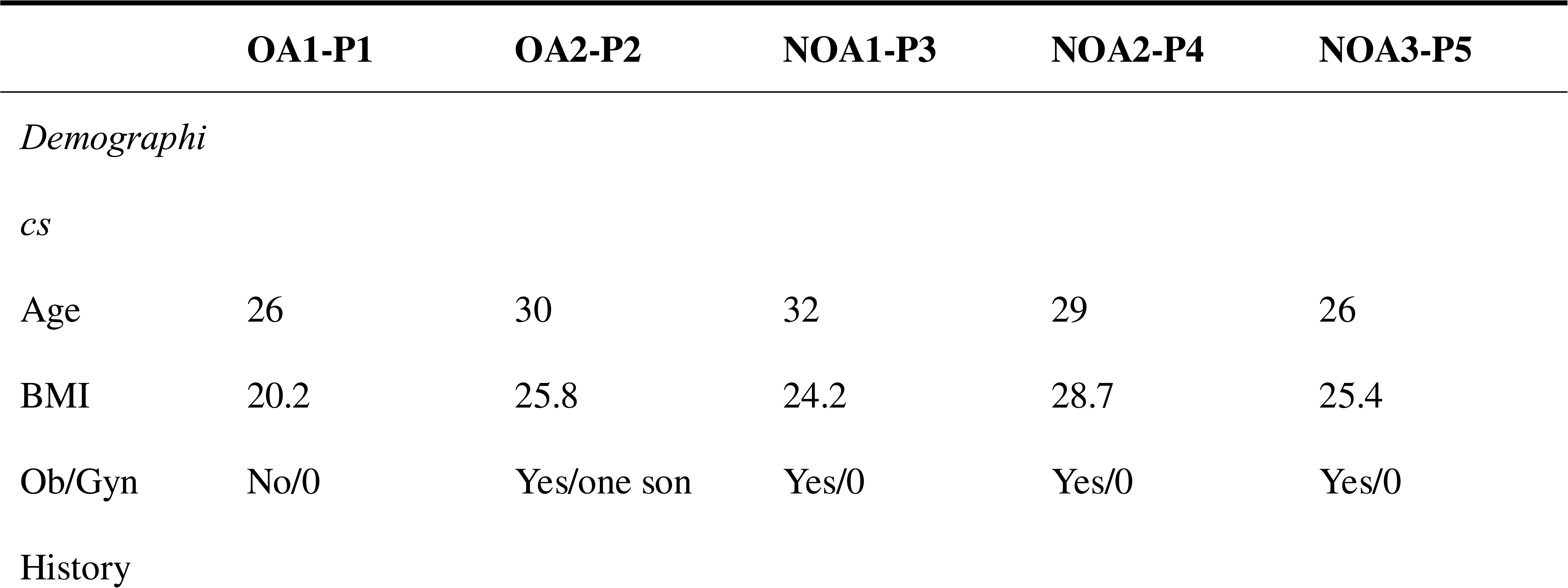
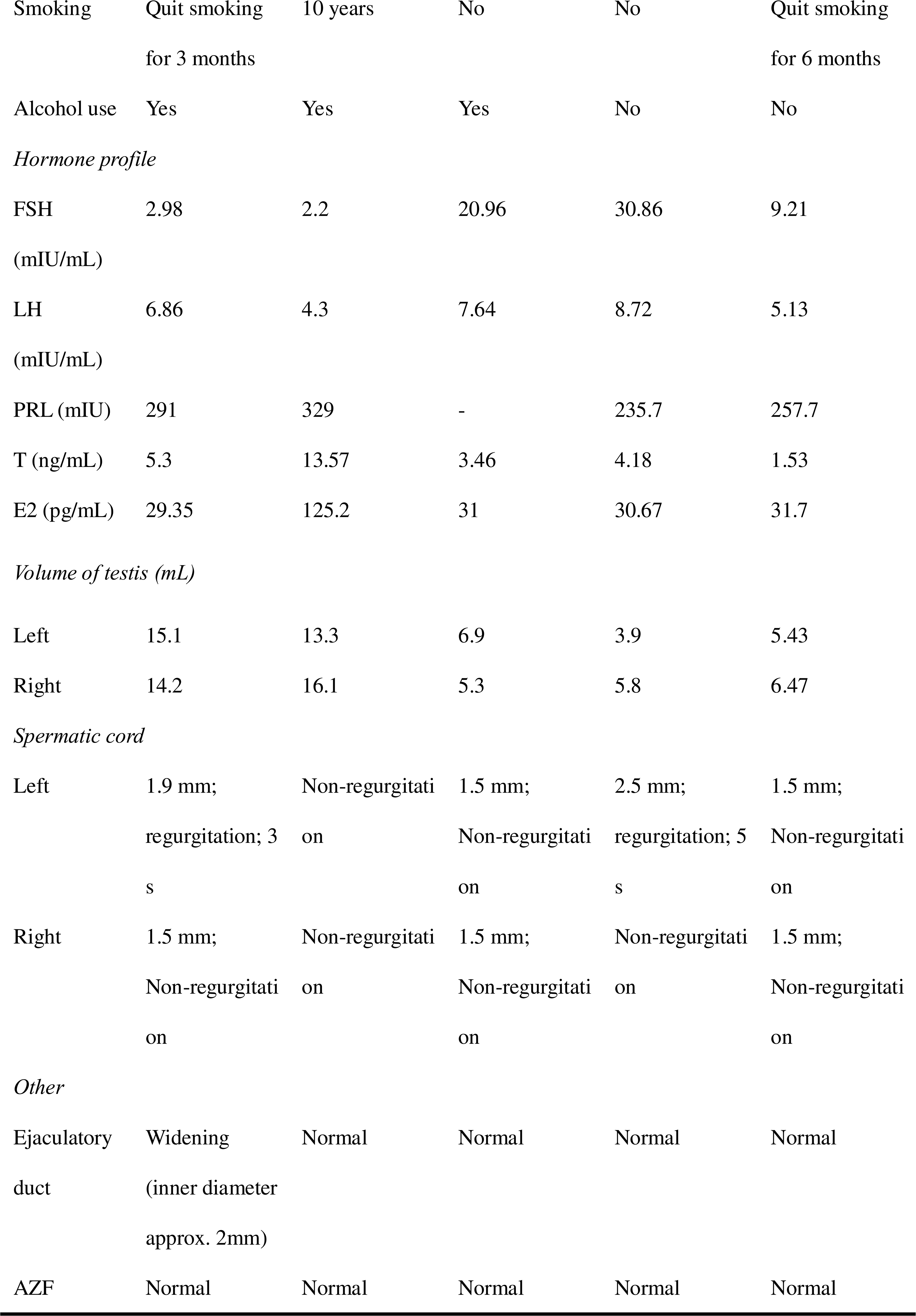
Patient characteristics in this study.

### Sample dissociation

Testicular tissues were washed three times in Dulbecco’s PBS (Gibco, Waltham, MA, USA) and the tunica albugineas of testes were removed. Testicular tissues were cut into small pieces of 1–2 mm and digested in 10 mL DMEM: F12 media (Gibco, Waltham, MA, USA) containing 1 mg/mL type IV collagenase for 15 min on a rotor at 37°C. Subsequently, tubules were washed again and incubated with 10 mL F12 media containing 500 μg/mL DNase I and 200 μg/mL trypsin for 15 minutes at 37°C. The digestion was stopped with the addition of fetal bovine serum (FBS) (Gibco, Waltham, MA, USA). The single cells were obtained by filtering through 40 μm cell-strainer. The cells were resuspended in 1 mL DPBS and 5 mL red blood cell lysis buffer to remove red blood cells. After that, the cells were washed with DPBS and cryopreserved in DPBS with 1% FBS (Gibco, Waltham, MA, USA) until further use. For scRNA-seq and scATAC-seq assay, the cryopreserved cells were thawed and incubated with a Dead Cell Removal Kit (Miltenyi Biotec) to clear dead/stressed cells in accordance with manufacturer’s instruction.

### scRNA-seq library preparation

scRNA-seq was carried out using 10x Chromium Single Cell 3’ Reagent referring to manufacturer’s instruction. In brief, viable single cells were resuspended in PBS with 0.04% bovine serum albumin (BSA; Sigma) and counted using the hemocytometer. Suspensions containing 5000 to 8000 cells cells per sample were mixed with RT-PCR reaction and loaded into the Single Cell Chip B and processed through the 10x controller for droplet production. After that, in-drop lysis and reverse transcription occurs and mRNA transcripts from single cells were barcoded to determine the cell origin. Following reverse transcription, barcoded cDNAs were purified, amplified by 12 cycles of PCR, end-repaired, and ligated with Illumina adapters. The final libraries were sequenced on the Illumina Novaseq 6000 platform with paired end 150 bp sequencing. Each sample was sequenced with depth of 100 G of raw reads.

### scRNA-seq data processing and analysis

The scRNA-seq data produced by Illumina NovaSeq 6000 sequencing were processed and mapped to human reference genome hg38. After mapping, the outputs were processed using the Seurat package in R. To filter out low-quality cells, we referred to the following criteria: cells with few genes per cell (<50) or plenty of molecules per cell (>20,000); cells with over 30% of mitochondrial genes. Normalization was carried out in accordance with the package manual (https://satij alab.org/seura t/v3.1/). We identified cell clusters with Uniform Manifold Approximation and Projection (UMAP). The differentially expressed genes were analyzed by the FindMarkers function. Gene set enrichment analysis was carried out using clusterProfiler R package. Regulatory gene network analysis was carried out by SCENIC.

### Pseudo-time analysis

We applied the Single-Cell Trajectories analysis utilizing Monocle2 (http://cole-trapnell-lab.github.io/monocle-release) using DDR-Tree and default parameter. Before Monocle analysis, we select marker genes of the Seurat clustering result and raw expression counts of the cell passed filtering. Based on the pseudo-time analysis, branch expression analysis modeling (BEAM Analysis) was applied for branch fate determined gene analysis.

### Velocity analysis

RNA velocity analysis was conducted using scVelo’s (version 0.2.1) generalized dynamical model. The spliced and unspliced mRNA was quantified by Velocity (version 0.17.17).

### Cell cycle analysis

To quantify the cell cycle phases for individual cell, we employed the CellCycleScoring function from the Seurat package. This function computes cell cycle scores using established marker genes for cell cycle phases as described in a previous study by Nestorowa et al. (2016). Cells showing a strong expression of G2/M-phase or S-phase markers were designated as G2/M-phase or S-phase cells, respectively. Cells that did not exhibit significant expression of markers from either category were classified as G1-phase cells.

### Cell communication analysis

To enable a systematic analysis of cell–cell communication molecules, we applied cell communication analysis based on the CellPhoneDB, a public repository of ligands, receptors and their interactions. Membrane, secreted and peripheral proteins of the cluster of different time point was annotated. Significant mean and Cell Communication significance (p-value<0.05) was calculated based on the interaction and the normalized cell matrix achieved by Seurat Normalization.

### SCENIC analysis

To assess transcription factor regulation strength, we applied the Single-cell regulatory network inference and clustering (pySCENIC, v0.9.5) workflow, using the 20-thousand motifs database for RcisTarget and GRNboost.

### scATAC-seq library preparation

scATAC-seq was conducted using 10x Chromium Single Cell ATAC Reagent (V1.1 chemistry) in accordance with manufacturer’s instruction. In short, cells were resuspended in PBS with 0.04% BSA (Sigma) and counted. Next, cells were cultured with cold lysis buffer on ice for 5 min. Tn5 transposase reaction was performed using the Tagment DNA Enzyme 1 (Illumina) at 37 °C for 30 min. Cells were processed for targeting a recovery of 10,000 cells per sample and amplification were performed with 12 cycles of PCR for library construction. The libraries were sequenced on Illumina Novaseq 6000 platform and each sample was sequenced with depth of 200M raw reads.

### scATAC-seq data preprocessing and analysis

The scATAC-seq data analysis was performed as previously described. In brief, the sequence reads were mapped to the human reference genome hg38. The R package Signac was used to process the generated outputs. The Latent Semantic indexing algorithm, T-distributed Stochastic Neighbor Embedding algorithm and Uniform Manifold Approximation algorithm are used for dimension reduction and information display. Each Motif of the cell can be annotated by associating identified open chromatin sites with the JASPAR database. By using CHIPseeker, the distribution of peak in different functional regions of the genome was annotated and statistically analyzed. Prediction of results for each cell type in scATAC-seq can be achieved by combining scRNA-seq data with scATAC-seq using KNN.

### Integrating scATAC-seq and scRNA-seq analysis

We primarily utilized Seurat for scRNA-seq data processing and Signac (version 1.13.0) for scATAC-seq data analysis. To establish connections between scRNA-seq and scATAC-seq datasets, we approximated gene transcriptional activity by measuring ATAC-seq signals within 2 kb upstream and the gene body regions using Signac’s GeneActivity function. These gene activity scores from scATAC-seq data, combined with gene expression levels from scRNA-seq, were input for canonical correlation analysis. Subsequently, we pinpointed anchors between the two datasets with the FindTransferAnchors function. We proceed to annotate scATAC-seq cells. For joint visualization in a UMAP plot, we projected RNA expression onto scATAC-seq cells using the pre-established anchors and integrated the datasets.

### Immunofluorescence

The testicular tissues were fixed in 4% paraformaldehyde at room temperature for 30min, permeabilized with 0.5% Triton X-100, and blocked with 1% bovine serum albumin (GIBCO, USA). The samples were incubated overnight at 4 °C with primary antibodies, including ST3GAL4 (13546-1-AP, Proteintech), A2M (ab109422, Abcam), DDX4 (ab270534, abcam), ASB9 (22728, SAB), TEX19 (AF6319, R&D), TSSK6 (H00083983-M02, Novus), PRAP1 (11932-1-AP, Proteintech), BST2 (H00000684-B02P, Novus), SOX9 (ab185966, Abcam), CCDC62 (25981-1-AP, Proteintech),TF (ab185966, Abcam), SOX2 (ab92494, Abcam), SPATS1 (NBP2-31037, Novus), C9orf57 (orb156207, Biorbyt), TSC21 (NBP1-91727, Novus), BEND4 (24711-1-AP, Proteintech), and SMCP (NBP1-81252, Novus). After washing with PBS, the samples were incubated with a secondary antibody (Abcam) for 1 h at room temperature. DAPI (Cell Signaling, MA, USA) was used to stain cell nucleus. After washing 3 time with PBS, the sections were performed for photograph under a fluorescence microscope (Carl Zeiss, Oberkochen, Germany).

## Results

### scRNA-seq analysis of human testicular tissues

Testicular histology was examined by HE staining. In the OA1-P1 group, seminiferous tubules displayed normal structure with well-organized layers of germ cells at various stages of spermatogenesis, including mature spermatozoa. In the OA2-P2 group, germ cells and spermatozoa were present but show disorganized arrangement, with sloughed cells observed in the lumen. In the NOA1-P3 group, no spermatozoa were present, but a small number of spermatocytes and spermatogonia were visible, along with various stages of spermatogenesis. In the NOA2-P4 group, germ cells and spermatozoa were present but exhibit disordered arrangement, with sloughed cells in the lumen. In the NOA3-P5 group, only Sertoli cells were observed, indicating impaired spermatogenesis (Figure 1A). To characterize the diversity of testicular cells, we performed scRNA-seq on testicular samples from three NOA patients and two OA patients (Figure 1B). After removing low quality cells, we obtained 23889 cells for scRNA-seq (Table S1). Notably, the quantity of annotated germ cells in patients categorized as NOA1, NOA2, and NOA3 gradually decreased to zero (Table S1). This observation is in complete accordance with the clinical features exhibited by the three patients, providing evidence for the reliability of our scRNA-seq data. UMAP analysis of scRNA-seq revealed 12 major cell types, including germ cells, Leydig cells, Sertoli cells, endothelial cells, peritubular myoid cells (PMCs), smooth muscle, schwann cell, macrophage, mast cells, T cells, B cells, and plasma cells (Figure 1C). A total of 41708 high quality cells were obtained from scATAC-seq analysis (Table S2), and they were also annotated with B cells, endothelial cells, PMCs, schwann cell, smooth muscle, T cells, germ cells, Leydig cells, macrophage, and Sertoli cells, which was similar to the scRNA-seq results (Figure 1D). Heatmap of marker genes in scRNA-seq data revealed that cell heterogeneity was obvious (Figure 1E). The well-known cell type markers were utilized to determine the cell clusters (Figure 1F). For instance, we observed that sertoli cells, in addition to specifically expressing classical markers AMH and SOX9 (Fröjdman, Harley, Pelliniemi, & biology, 2000; Lasala et al., 2011), also exhibited strong expression of APOA1 (Figure 1F). Furthermore, we identified that the PMCs not only expressed the classical marker MYH11 (Lottrup et al., 2014), which is concurrently highly expressed in smooth muscle cells, displaying limited specificity, but also demonstrated higher specificity in the expression of DPEP1, suggesting its potential as a specific marker for PMCs. Similar to the scRNA-seq results, the activity of these markers was high in the corresponding cell subtypes (Figure S1A). Figure S1B showed the chromatin accessibility of these markers within the cells. Integrating scATAC-seq and scRNA-seq analysis revealed indicated substantial overlap between the cells profiled in the scRNA-seq and scATAC-seq data, implying that for most cell populations, shifts in chromatin accessibility were aligned with coordinated alterations in gene expression levels (Figure S2A). These findings indicate that the data from this study can serve for cellular classification.

**Figure 1.**
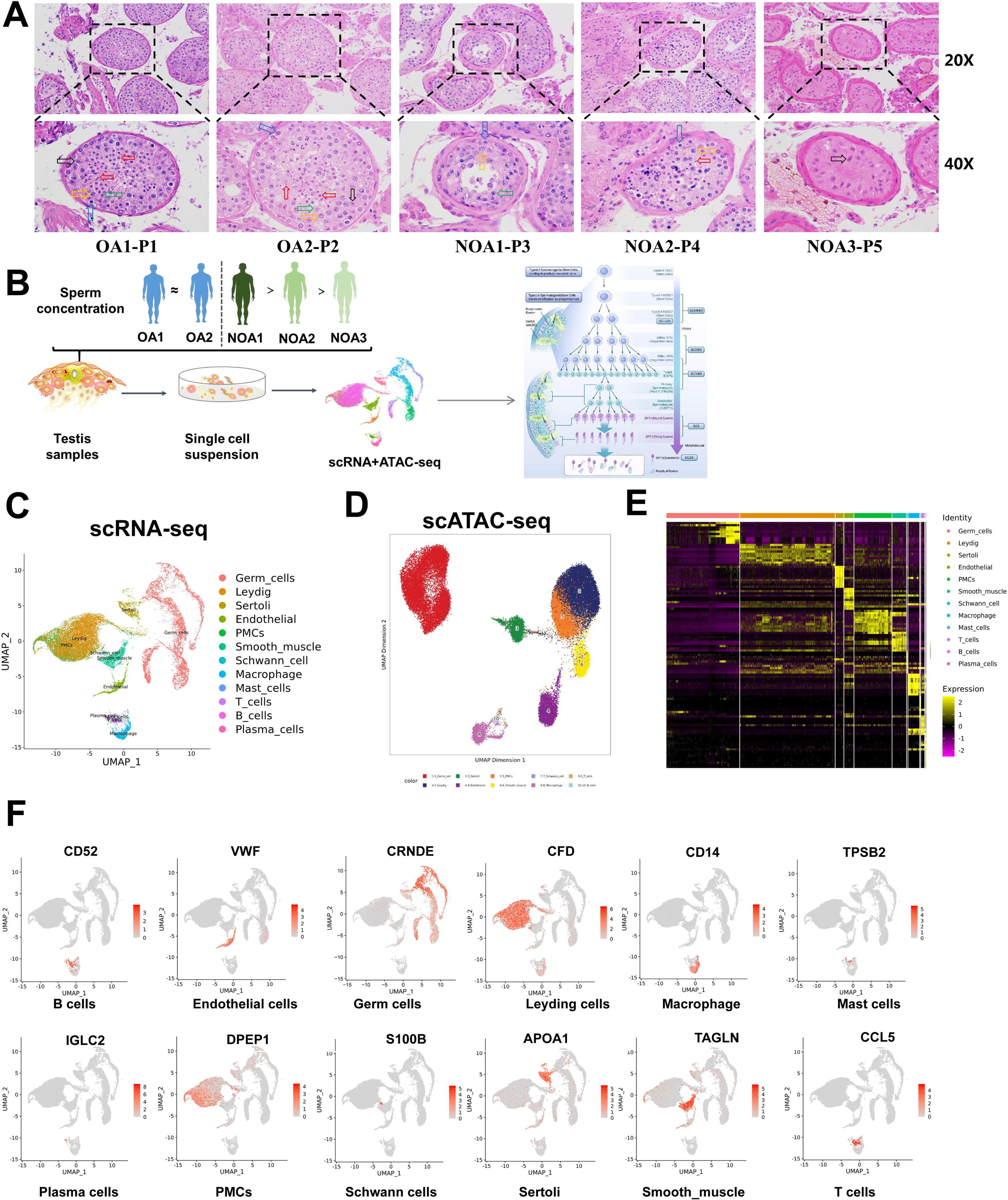
Overview of major cell types and cellular attributes of testes of OA and NOA patients. **(A)** Representative images of HE staining of testicular tissues of OA and NOA patients. (**B**) Schematic of the experimental design for scRNA-seq and scATAC-seq. (**C**) UMAP analysis of human testicular cells in scRNA-seq results. (**D**) UMAP analysis of human testicular cells in scATAC-seq results. (**E**) Heatmap of expression of markers for the 12 cell types. (**F**) UMAP analysis of testicular cell population.

### Germ cell subtypes reveal distinct molecular features

To explore the heterogeneity of germ cells during normal development, we re-clustered the germ cells and identified 12 subpopulations, including spermatogonial stem cell-0 (SSC0), SSC1, SSC2, and Diffing_SPG, Diffed_SPG, Pre-Leptotene, Leptotene_Zygotene, Pachytene, Diplotene, SPT1, SPT2, and SPT3 (Figure 2A). Integrating scATAC-seq and scRNA-seq analysis the distributions between the same cell types were very alike (Figure S2B). Heatmap of scRNA-seq results showed the the markers successfully distinguished between different germ cell subpopulations (Figure S3A). Violin plots showed germ cell subtype marker gene expression in different germ cells (Figure S3B). Heatmap of germ cell markers and motifs in scATAC-seq results were shown in Figure S4A and Figure S4B. Subsequently, we attempted to characterize the sequential developmental relationships among 12 germ cell subpopulations through pseudotime analysis, approaching this issue from a developmental perspective. As shown in Figure 2B, germ cells could be divided into three states, which was in accordance with the biological sequence of continuous transition of spermatogenesis. States 2 and 3 primarily consist of SSC-Diffed SPG cells, and these cells are not completely segregated in terms of the cell states, particularly with SSC0-SSC3 positioned at two initial branches without complete separation (Figure 2B, Figure S3C). This suggests a potential cyclical and self-renewing state that is not entirely independent between SSC1 and SSC2. Cells from the Pre-L to SPT3 stages all belong to state 1, following the developmental chronological order of sperm generation. Therefore, the phenomenon of distinct subpopulations existing independently while sharing markers with other subpopulations suggests that germ cells from different subtypes may not have fully differentiated into mature states during the course of evolution, remaining in an intermediate state of continuous differentiation.

**Figure 2.**
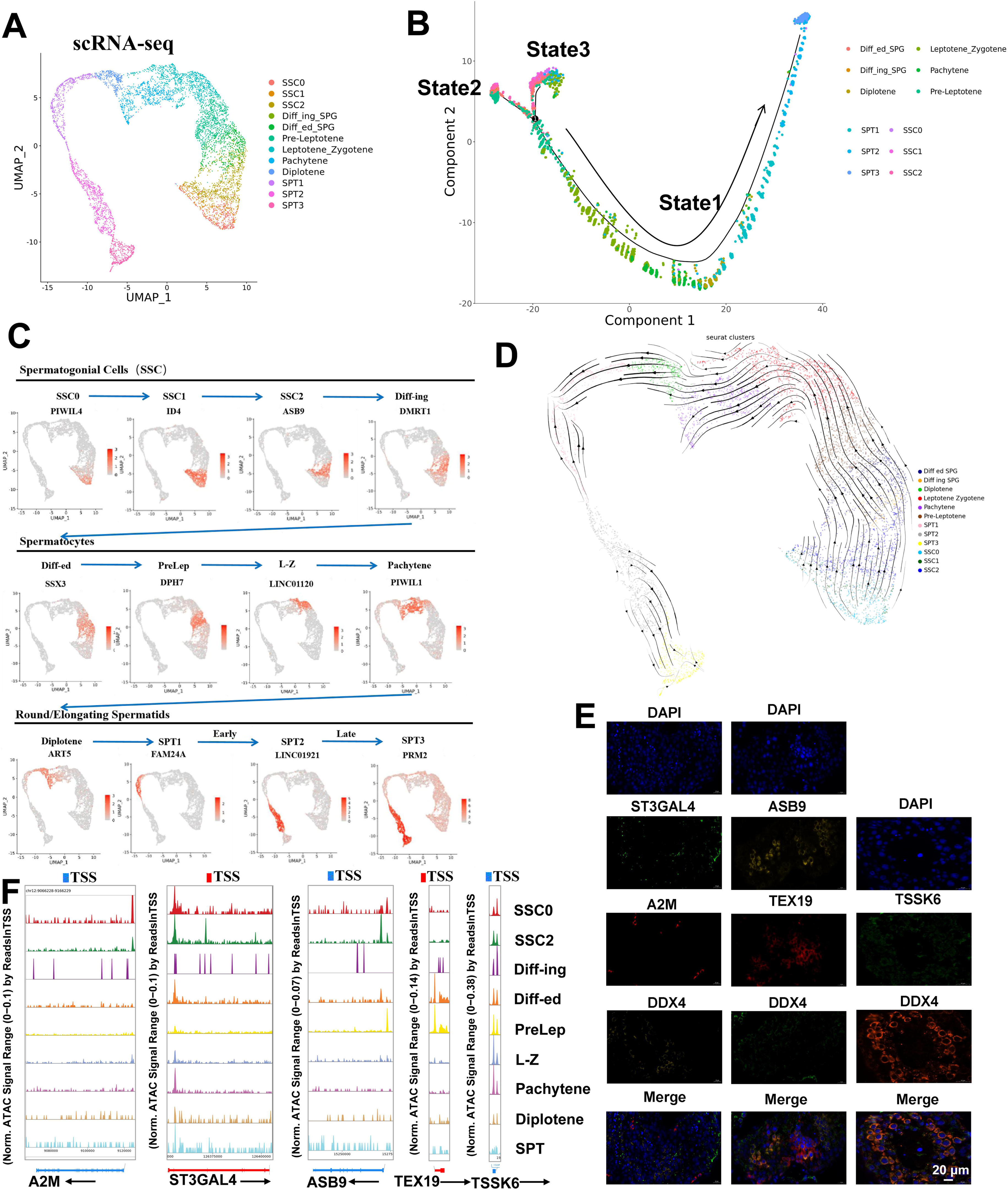
Identification of germ cell subtypes. (**A**) UMAP analysis of 12 germ cell subsets in scRNA-seq results. (**B**) Pseudotime analysis of 12 germ cell subpopulations. (**C**) Gene expression patterns of marker genes on UMAP plots. (**D**) State of germ cell subsets using pseudotime analysis. (**E**) Immunofluorescence double staining of markers of newly identified spermatocyte subpopulations with the classical marker DDX4. (**F**) Accessibility peak plots of markers for newly identified germ cell subpopulations.

Furthermore, we presented the expression distribution of markers for different subpopulations across three developmental stages in the chronological order of germ cell maturation (Figure 2C). Interestingly, the positions of different subpopulations of markers on UMAP show a trend of right-to-left panning as the arrows point to the temporal developmental order of sperm cells, which was supported by RNA velocity stream UMAP (Figure 2D). Among them, SPT1 cells can be clearly distinguished from SPT2-3 cells using FAM24A, but between SPT2 and SPT3 cells, there is no precise and specific marker that completely segregates these two cell types. We speculate that this lack of differentiation may be due to the intense morphological changes occurring in the sperm cells during this period, resulting in relatively minor differences in gene expression.

Finally, to further validate the reliability of the identified new potential markers of germ cells, we co-stained them with a classical marker DDX4 using immunofluorescence. We discovered that ST3GAL4, A2M, and DDX4 were co-expressed in testes tissues. ASB9 and TEX19 were also co-expressed with DDX4 were in testes tissues. TSSK6 exhibited similar expression pattern to that of DDX4 in testes tissues (Figure 2E). In addition, closely resembled their expression patterns in different subpopulations, we found that the region transcription start/promoter site to A2M and ASB9 was accessible in the SSC0, SSC2, and Diffing_SPG but was not accessible in other cell subtypes, with much higher levels of openness (Figure 2F). The region transcription start site to TEX19 was accessible in the Diffed_SPG but was not accessible in other cell subtypes, which also closely resembled TEX19 expression pattern. Collectively, these results suggest that these new markers localize to testicular tissue and have the same ability to label sperm cells as DDX4.

### Cell cycle and transcriptional dynamics analysis of germ cell differentiation

Most adult tissues are sustained by resident adult stem cells responsible for maintaining tissue function and integrity. As aging or damaged cells undergo apoptosis, resident adult stem cells are activated to generate new cells. Within various tissues, a dichotomy exists between rapidly cycling (involved in tissue repair, EOMES - GFRA1^+^) and quiescent stem cells (reserve cells, EOMES^+^GFRA1^+^) (Sharma et al., 2018). To investigate the cellular developmental states of distinct sperm subpopulations, we conducted an analysis of the cell cycle in sperm cells. As shown in Figure 3A and 3B, SSC0/1 and SSC2 were quiescent stem cells and rapidly cycling cells, respectively. Furthermore, to explore the biological function of germ cells, we used transcriptional dynamics to analyze the important events and time points in the whole transcriptional process of spermatogenesis. Maintenance of SSC pluripotency, whether in the G0/G1 phase and G2/M arrest SSC0/SSC1, or rapid cycling SSC2, are transcriptionally governed by the downregulation of ID4 (Figure 3C). The pivotal driving genes promoting the transition of SSC2 into differentiating Diffing SPG cells and progressing through mitosis are upregulated DMRT1 and PABPC4 (Figure 3C). As BEND2 takes the central regulatory position, germ cell development officially proceeds into meiosis, marking the transitioned cells (Diffed SPG) in a mixed transitional state concurrently undergoing both mitosis and meiosis (Figure 3C). The molecular markers distinguishing between the PreLep and SPT3 subpopulations included C18orf63, MNS1, CCDC42, LDHC, TMIGD3, AC113189.2, and FLJ40194 (Figure 3C). Results showed that there is no distinctly discernible regulatory factor between PreLep and L-Z, with C18orf63 being a crucial differentiator, upregulated in PreLep and downregulated in L-Z. Finally, we summarized the spermatogenesis at each stage and major events in the cell cycle (Figure 3D).

**Figure 3.**
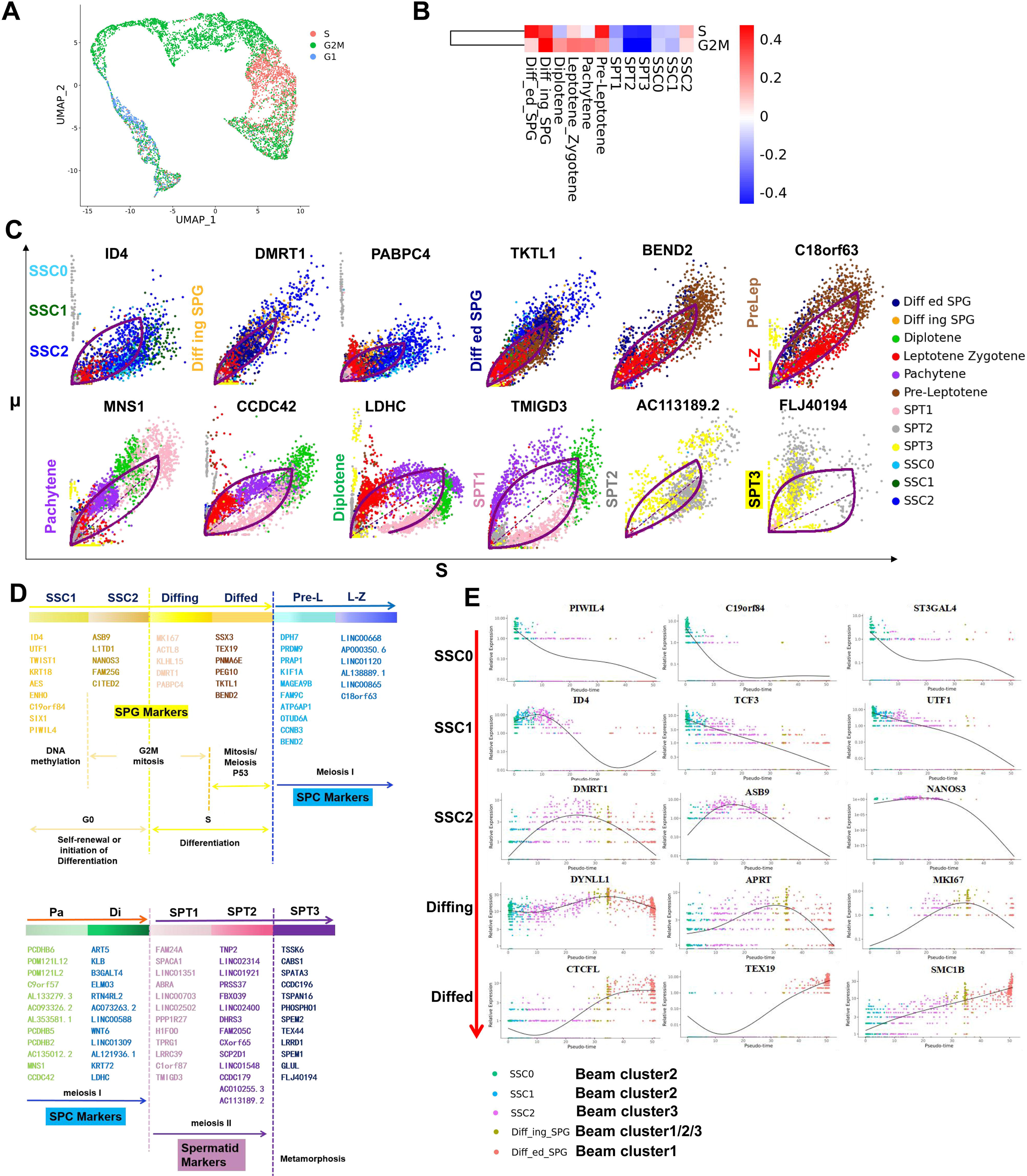
Analysis of germ cells at different stages. (**A**) Germ cell subsets at different stages of the cell cycle. (**B**) Heatmap of cell cycle in germ cells. (**C**) Transcriptional dynamics of driver genes of germ cell subtypes. (**D**) The main stage of spermatogenesis. (**E**) Beam analysis of differentially expressed genes at different stages of SSC.

Subsequently, Beam analysis was performed to unveil the critical transcription factors involved in sperm differentiation. Figure S5A hierarchically clustered the top 50 transcription factors into three groups, presenting an expression heatmap of fate-determining genes associated with the differentiation from SSC0 to SPG state. The expression pattern of transcription factors within cluster 1 gradually increased along the temporal trajectory (Figure S5B), predominantly involved in cytoplasmic translation GO functions (Figure S5C); Cluster 2 exhibited an initial increase followed by a decline (Figure S5B), governing positive regulation of translation and regulation of G2M transition of mitotic cell cycle (Figure S5C); transcription factors within cluster 3 showed a sustained decrease in expression (Figure S5B) and were enriched in cell cycle and meiotic cell cycle-related GO terms (Figure S5C). As shown in Figure 3E, PIWIL4, C19orf84 and ST3GAL4 were specifically expressed in SSC0 (belong to cluster 2). ID4, TCF3 and UTF1 were specifically expressed in SSC1 (belong to cluster 2). DMRT1, ASB9 and NANOS3 were the main specifically expressed genes in SSC2 (belong to cluster 3). DYNLL1, APRT and MKI67 were specifically expressed in Diffing-SPG (belong to cluster 1/2/3). CTCFL, TEX19 and SMC1B were specifically expressed in Diffed-SPG (belong to cluster 1). Heatmaps of transcription factors showed that the transcription of SSC1 and SSC2 was very active during reproductive development (Figure S5D). In contrast, SPT2/3 was almost transcribed in silence. Diffing/ed served as a connecting link between the preceding and the following, but its transcriptional state was closer to SSC (Figure S5D). In conclusion, we obtained 50 transcription factors that are closely related to sperm differentiation.

### Sertoli cells from human testicular tissues are composed of heterogeneity population

To explore the heterogeneity of Sertoli cells, we re-clustered the Sertoli cells and identified 8 subclusters (Figure 4A). Integrating scATAC-seq and scRNA-seq analysis the distributions between the same cell types were very alike (Figure S2C). The top differentially expressed genes were utilized to determine the Sertoli cell clusters using heatmap (Figure 4B) and violin plots (Figure 4C). Of note, no cluster-specific distinctive features were identified in SC1, as the marker genes (TF, SLC16A1 and CCNL1) of SC1 were also highly expressed in other Sertoli cells subclusters (Figure 4C). SOX2, NF1B and COL27A1 were preferentially expressed in SC2. SPATS1, LINC01120 and LINC01206 were specifically expressed in SC3. BEND2, PRAP1 and TEX1 were specifically expressed in SC4. PLPP2, BEND4 and RHOXF1 were specifically expressed in SC5. MLC1, SMCP and NUPR2 were highly expressed in SC6. FLRT2, CFH and BST2 were specifically expressed in SC7. TMIGD3, PGAM2 and CCDC62 were specifically expressed in SC8.

**Figure 4.**
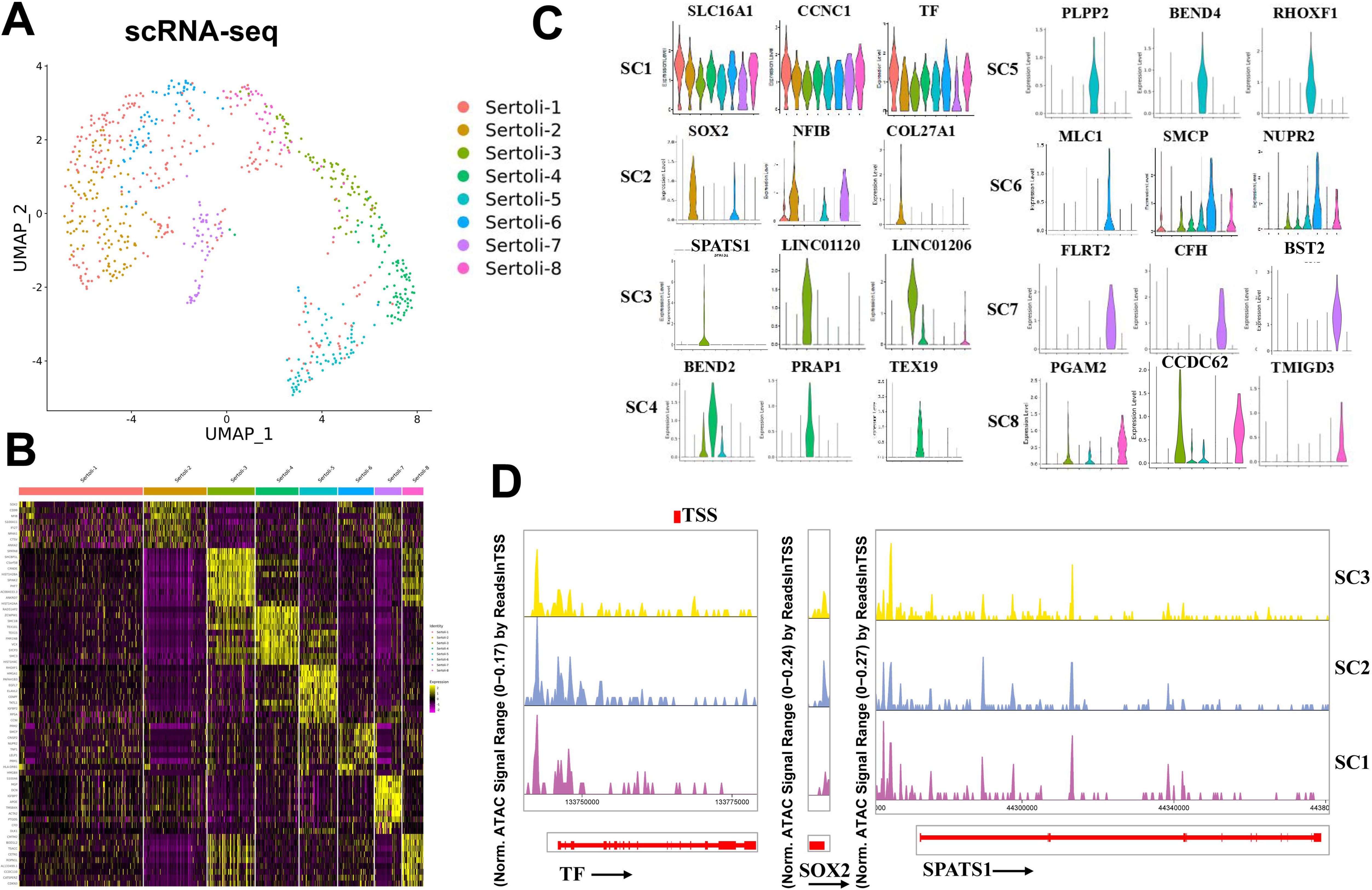
Identification of Sertoli cell subtypes and epigenetic analysis. (**A**) UMAP analysis showed 8 Sertoli cell subsets in scRNA-seq results. (**B**) Heatmap of markers of differentially expressed genes in eight Sertoli cell subsets. (**C**) Gene expression patterns of Sertoli cell subtype marker genes on violin plots. (**D**) Accessibility peak plots of markers for newly identified Sertoli cell subpopulations.

Cell differentiation is accompanied by the expression of genes controlled by cis-regulatory elements, which must be in an accessible state to function properly. Therefore, scATAC-seq analysis was performed on the same Sertoli cells as those used in scRNA-seq analysis, and a chromatin accessibility landscape for individual cell marker was delineated (Figure 4D). Because the number of cells in the SC4-SC8 subpopulation was too small to calculate a peak plot, only the peak plots of markers in the SC1-SC3 subpopulations are displayed. The promoter regions of TF were all highly accessible in the SC1-SC3 subpopulations, and relatively higher openness was observed for SOX2 in the SC2 subgroup. In addition to the high openness we observed in the promoter region of the SPATS1, there were also multiple accessible regions within the gene in the SC1-SC3 subpopulations.

### Developmental trajectories of Sertoli cells and identified three novel subpopulations

To analyze the origin and maturation process of Sertoli cells, pseudotime trajectory analysis was carried out. The trajectory was determined to initiate with State 3 as beginning and reached a terminally differentiate state of State 2 and State 1 (Figure 5A). According to Figure 5B, State 1 mainly included SC3/4/5 clusters, State 2 mainly included SC2/6/8 clusters, State 3 mainly included SC1/7 clusters. Figure 5C demonstrated RNA velocity heatmap of dynamic evolutionary Sertoli cells on the three States. Although most studies have categorized the development of Sertoli cells after birth into two stages: immature and mature, the existence of an intermediate or transitional state remains largely unknown. However, in this study, we identified three distinct states of Sertoli cells, with State 1 expressing markers EGR3, CTSL, PCNA, MK167, and KRT18 associated with immature Sertoli cells (Zhao et al., 2020), while State 3 expressed markers, such as TF, HOPX, and DEF119, indicative of mature Sertoli cells (Zhao et al., 2020). Consequently, we defined State 1 and State 3 as immature and mature Sertoli cells, respectively. State 2, falling between immature and mature states, does not belong to either category. Therefore, it is defined as a further maturation Sertoli cells, aligning with previously reported findings in the literature (Guo et al., 2021; Zhao et al., 2020). According to the latent time of RNA rate, the differentiation process is SC3, SC4, SC5, SC2, SC6, SC8, SC1 and SC7 (Figure 5D). Based on the above result, we identified three novel Sertoli cell subtypes (namely SC4/7/8) and their specific markers, and defined SC4 (PRAP1) as immature, SC7 (BST2) as mature, and SC8 (CCDC62) as further mature (Figure 5D). To validate these novel identified markers in Sertoli cells, we co-stained them with SOX9, a classical marker of Sertoli cells. As shown in Figure 5E, we discovered that PRAP1, BST2, and CCDC62 were co-expressed with SOX9 in testes tissues.

**Figure 5.**
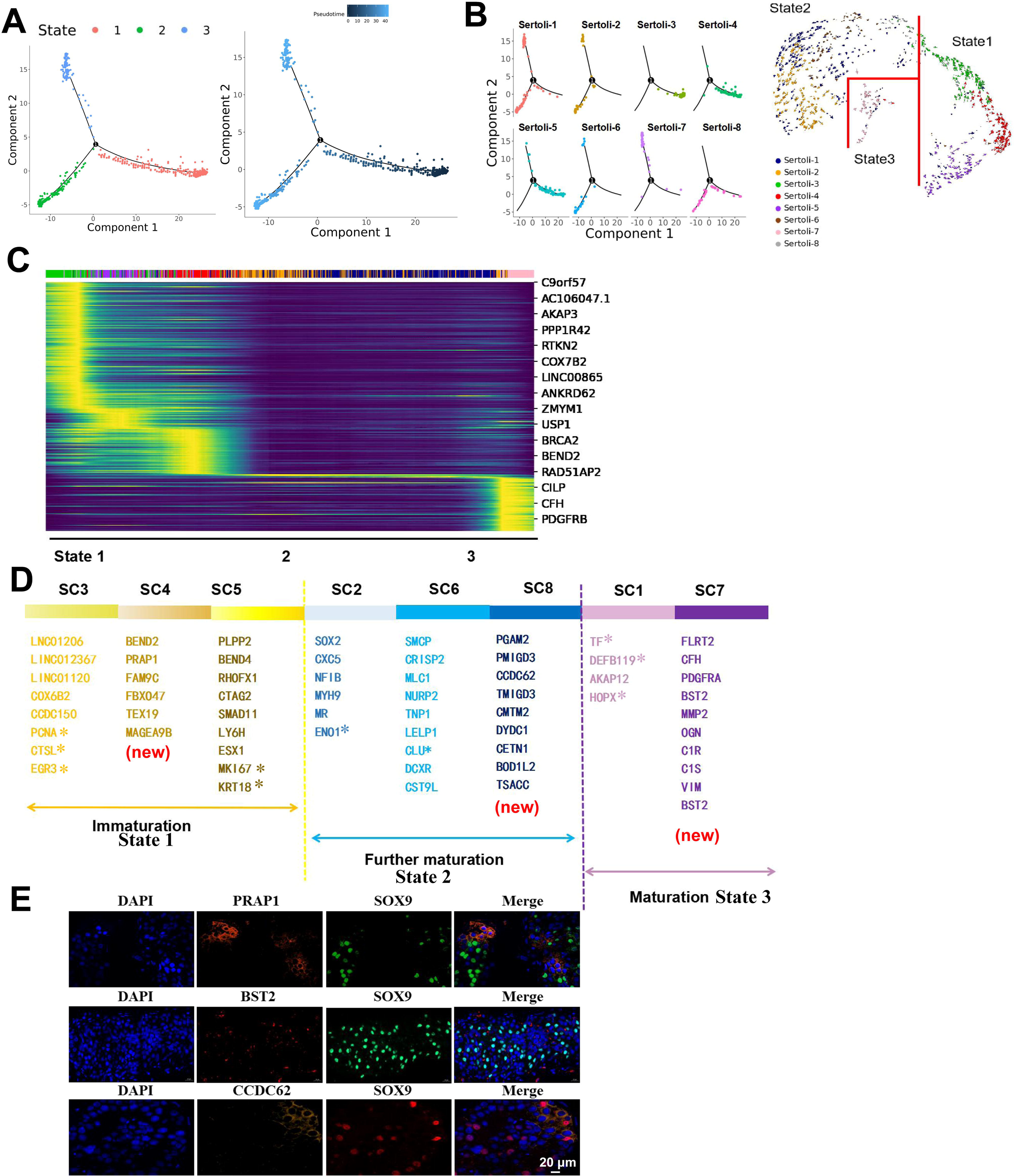
The development of Sertoli cells. (**A**) State of Sertoli cells using pseudotime analysis. (**B**) Distribution of Sertoli cell subtypes in different states. (**C**) Potential temporal differentiation of SC subgroups using RNA velocity. (**D**) Sertoli cells division diagram and representative markers. (**E**) Immunofluorescence of Sertoli cell subtype markers in testicular tissues from OA patients. The scale bar represents 20 μm.

GO analysis of Sertoli cells in different states was shown in Figure S5E. Sertoli cells in state 1 were mainly involved in cell cycle, meiotic cell cycle and spermatogenesis. Sertoli cells in state 2 were mainly involved in platelet degranulation, lipid metabolic process and steroid biosynthetic process. Sertoli cells in state 3 were mainly involved in extracellular matrix organization, SRP-dependent cotranslational protein targeting to membrane and viral transcription. Further Qusage analysis was performed for Sertoli cells (Figure S5F). Sertoli cells in state 1 was mainly involved in Cell cycle, DNA replication, and Mismatch repair. Sertoli cells in state 2 was involved in D−Glutamine and D−glutamate metabolism, Ribosome, and Valine, leucine and isoleucine biosynthesis. Sertoli cells in state 3 participated in ECM−receptor interaction, and Protein digestion and absorption. Overall, Sertoli cells performed different functions at different stages.

### The existence of Sertoli cell subtypes is more crucial for spermatogenesis than its quantity

The highly interdependent structural relationship between Sertoli cells and germ cells has long been considered evidence of their close functional association (Griswold, 1995). For instance, some germ cells, due to their structural affinity with Sertoli cells, cannot be fully separated, when cultured independently in vitro, these germ cells exhibit very short survival times, however, the addition of Sertoli cells or the use of Sertoli cells-conditioned medium significantly improves the survival of germ cells (La et al., 2018; Mohammadi-Sardoo et al., 2021; Risley & Morse-Gaudio, 1992). To determine which subtype of Sertoli cells is more closely involved in spermatogenesis, we analyzed spermatogenesis and Sertoli cells distribution in NOA (experimental group) and OA (control group) groups (Figure 6). The number proportion of germ cell subsets in NOA1/2 and OA1/2 was shown in Figure 6A. The number of germ cells in NOA3 patients was 0. The proportion of Sertoli cell subsets in NOA1/2/3 and OA1/2 was shown in Figure 6B. It can be seen that the number of Sertoli cells in NOA2 samples is higher than that in NOA3 samples, but the number of Sertoli cells in three NOA samples were lower than that in OA samples, indicating that the number of Sertoli cells is somewhat correlated with spermatogenesis.

**Figure 6.**
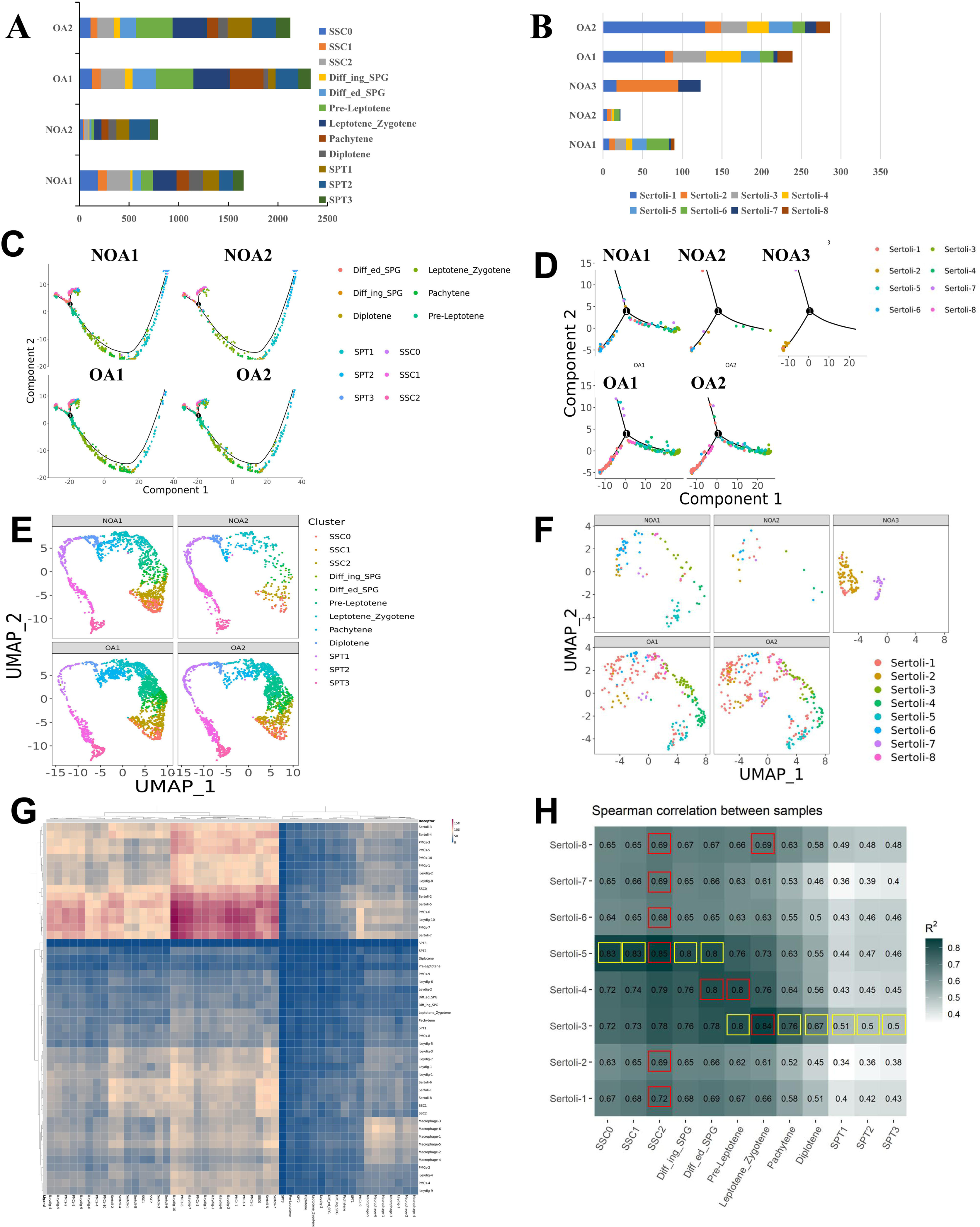
Correlation analysis of germ and Sertoli cell expression in different groups. (**A**) Cell proportion of germ cell subtypes in different groups. (**B**) Cell proportion of Sertoli cell subtypes in different groups. (**B**) Pseudotime analysis of germ cell subtypes in different groups. (**D**) Pseudotime analysis of Sertoli cell subtypes in different groups. (**E**) Distribution map of germ cell subtypes in different samples. (**F**) Distribution map of Sertoli cell subtypes in different samples. (**G**) The interaction between germ cells and other cells. (**H**) Spearman correlation between germ and Sertoli cells.

Moreover, interestingly, we observed a complete absence of immature Sertoli cells, especially SC3, in the testicular tissue of patients with NOA3 who exhibited a total absence of sperm, with only a small population of mature SC7 cells present (Figure 6C and 6D), suggesting that the absence of sperm in NOA3 patients may be associated with Sertoli cells SC3. For NOA2 samples, although the number of Sertoli cells was less than that of NOA3, SC3 was not missing in NOA2, so spermatogenesis was only partially affected (Figure 6C – 6F). In conclusion, these data suggested that whether or not the critical subtype SC3 is missing is more important for spermatogenesis than the number of Sertoli cells.

### Co-localization of subpopulations of Sertoli cells and germ cells

To determine the interaction between Sertoli cells and spermatogenesis, we applied Cell-PhoneDB to infer cellular interactions according to ligand-receptor signalling database. As shown in Figure 6G, compared with other cell types, germ cells were mainly interacted with Sertoli cells. We futher performed Spearman correlation analysis to determine the relationship between Sertoli cells and germ cells. As shown in Figure 6H, State 1 SC3/4/5 were tended to be associated with PreLep, SSC0/1/2, and Diffing and Diffed-SPG sperm cells (R > 0.72). Interestingly, SC3 was significantly positively correlated with all sperm subpopulations (R > 0.5), suggesting an important role for SC3 in spermatogenesis and that SC3 is involved in the entire process of spermatogenesis. Subsequently, to understand whether the functions of germ cells and Sertoli cells correspond to each other, GO term enrichment analysis of germ cells and sertoli cells was carried out (Figure S6, S7). We found that the functions could be divided into 8 categories, namely, material energy metabolism, cell cycle activity, the final stage of sperm cell formation, chemical reaction, signal communication, cell adhesion and migration, stem cells and sex differentiation activity, and stress reaction. These different events were labeled with different colors in order to quickly capture the important events occurring in the cells at each stage. As shown in Figure S6, we discovered that SSC0/1/2 was involved in SRP-dependent cotranslational protein targeting to membrane, and cytoplasmic translation; Diffing SPG was involved in cell division and cell cycle; Diffied SPG was involved in cell cycle and RNA splicing; Pre-Leptotene was involved in cell cycle and meiotic cell cycle; Leptotene_Zygotene was involved in cell cycle and meiotic cell cycle; Pachytene was involved in cilium assembly and spermatogenesis; Diplotene was involved in spermatogenesis and cilium assembly; SPT1 was involved in cilium assembly and flagellated sperm motility; SPT2 was involved in spermatid development and flagellated sperm motility; SPT3 was involved in spermatid development and spermatogenesis. As shown in Figure S7, SC1 were mainly involved in cell differentiation, cell adhesion and cell communication; SC2 were involved in cell migration, and cell adhesion; SC3 were involved in spermatogenesis, and meiotic cell cycle; SC4 were involved in meiotic cell cycle, and positive regulation of stem cell proliferation; SC5 were involved in cell cycle, and cell division; SC6 were involved in obsolete oxidation−reduction process, and glutathione derivative biosynthetic process; SC7 were involved in viral transcription and translational initiation; SC8 were involved in spermatogenesis and sperm capacitation. The above analysis indicated that the functions of 8 Sertoli cell subtypes and 12 germ cell subtypes were closely related.

To further verify that Sertoli cell subtypes have “stage specificity” for each stage of sperm development, we firstly performed HE staining using testicular tissues from OA3-P6, NOA4-P7 and NOA5-P8 samples. The results showed that the OA3-P6 group showed some sperm, with reduced spermatogenesis, thickened basement membranes, and a high number of sertoli cells without spermatogenic cells. The NOA4-P7 group had no sperm initially, but a few malformed sperm were observed after sampling, leading to the removal of affected seminiferous tubules. The NOA5-P8 group showed no sperm in situ (Figure 7A). Immunofluorescence staining in Figure 7B was performed using these tissues for validation. ASB9 (SSC2) was primarily expressed in a wreath-like pattern around the basement membrane of testicular tissue, particularly in the OA group, while ASB9 was barely detectable in the NOA group. SOX2 (SC2) was scattered around SSC2 (ASB9), with nuclear staining, while TF (SC1) expression was not prominent. In NOA patients, SPATS1 (SC3) expression was significantly reduced. C9orf57 (Pa) showed nuclear expression in testicular tissues, primarily extending along the basement membrane toward the spermatogenic center, and was positioned closer to the center than DDX4, suggesting its involvement in germ cell development or differentiation. BEND4, identified as a marker fo SC5, showed a developmental trajectory from the basement membrane toward the spermatogenic center. ST3GAL4 was expressed in the nucleus, forming a circular pattern around the basement membrane, similar to A2M (SSC1), though A2M was more concentrated around the outer edge of the basement membrane, creating a more distinct wreath-like arrangement. In cases of impaired spermatogenesis, this arrangement becomes disorganized and loses its original structure. SMCP (SC6) was concentrated in the midpiece region of the bright blue sperm cell tail. In the OA group, SSC1 (A2M) was sparsely arranged in a rosette pattern around the basement membrane, but in the NOA group, it appeared more scattered. SSC2 (ASB9) expression was not prominent. BST2 (SC7) was a transmembrane protein primarily localized on the cell membrane. In the OA group, A2M (SSC1) was distinctly arranged in a wreath-like pattern around the basement membrane, with expression levels significantly higher than ASB9 (SSC2). TSSK6 (SPT3) was primarily expressed in OA3-P6, while CCDC62 (SC8) was more abundantly expressed in NOA4-P7, with ASB9 (SCC2) showing minimal expression. Taken together, germ cells of a particular stage tended to co-localize with Sertoli cells of the corresponding stages. Germ cells and sertoli cells at each differentiation stage were functionally heterogeneous and stage-specific (Figure 8). This suggests that each stage of sperm development requires the assistance of sertoli cells to complete the corresponding stage of sperm development.

**Figure 7.**
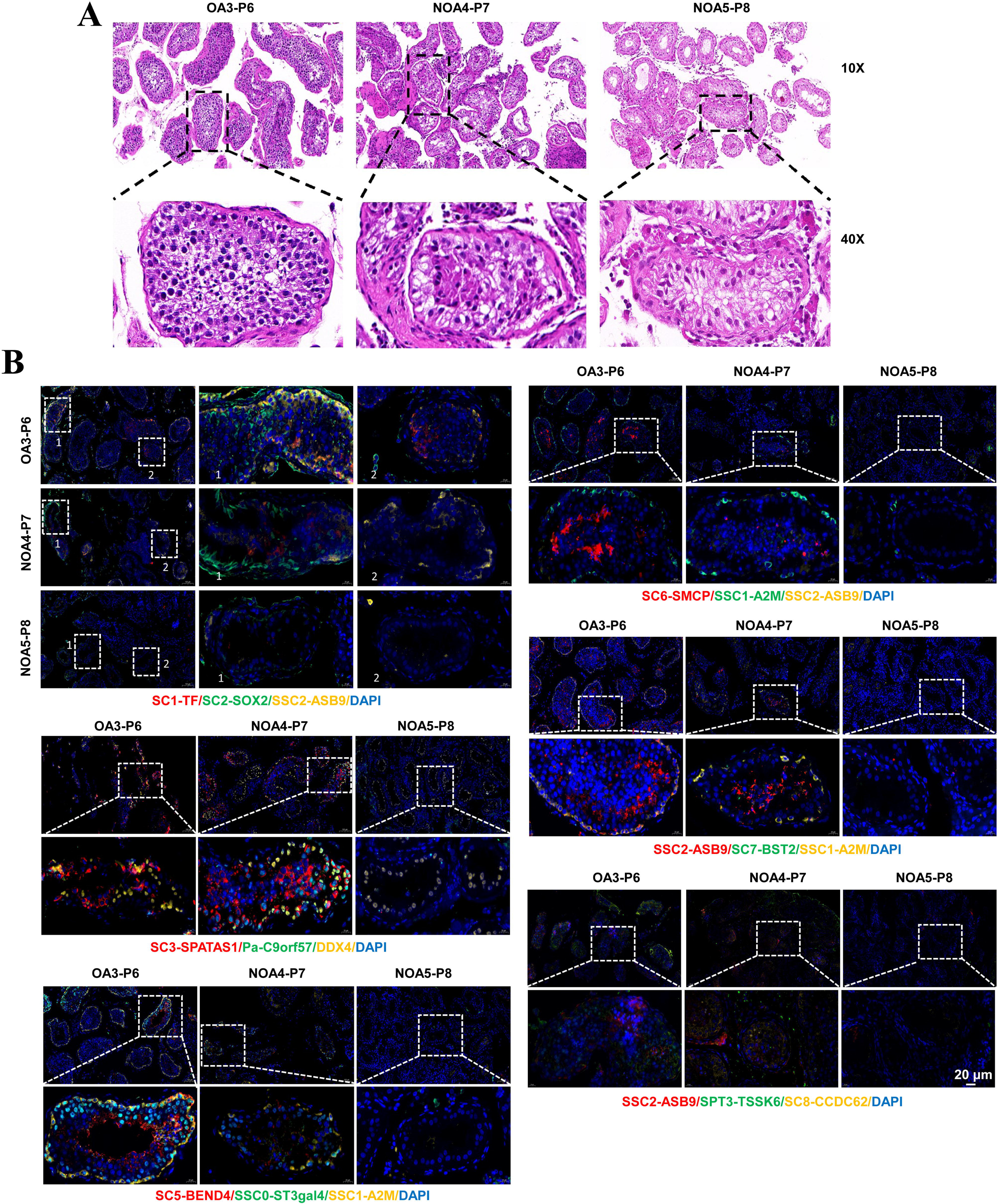
Co-localization of subpopulations of Sertoli cells and germ cells. (A) Representative images of HE staining of testicular tissues of OA and NOA patients. (B) Immunofluorescence analysis of germ cell and Sertoli cell subtype markers in testicular tissues. Blue DAPI indicates nuclei. The colors of the markers and cells indicate the fluorescent color of the corresponding antibody. The scale bar represents 20 μm.

**Figure 8.**
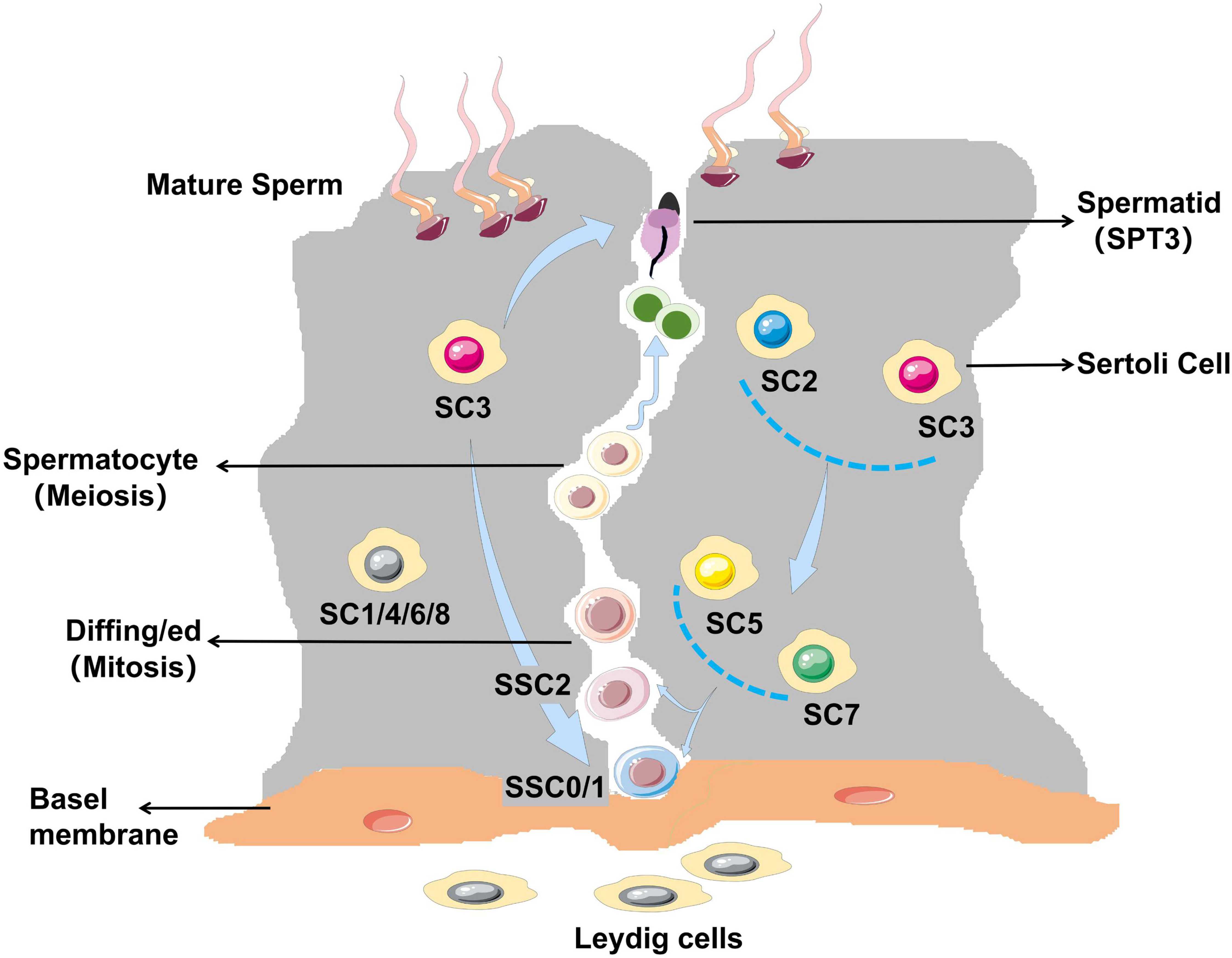
Schematic diagram of spermatogenesis and stage specificity of sertoli cell subtypes.

### Notch1/2/3 signaling participate in the interaction between Sertoli cells and germ cells

To explore the mechanism of interaction between Sertoli cells and germ cells, we conducted Cell Phone analysis. Results showed that Notch1/2/3 signaling were involved in the interaction between Sertoli cells and germ cells (Figure S8A). Interestingly, a chromatin accessibility landscape in scATAC-seq data showed that promoter regions of Notch1 only accessible in the SC1-SC3 subpopulations not in germ cells (Figure S8B), suggesting that Notch1 may be expressed by germ cells. The openness of promoter region of Notch2 was observed not only in germ cells but also in SSC2 and Pachytene celltypes. Notch3 accessibility was only observed within the gene in different cell subtypes. Overall, Notch1/2/3 signaling are involved in the interaction between Sertoli cells and germ cells.

### Leydig cell heterogeneity and development

Leydig cells are the main source of androgens and exist in the interstitial space between the seminiferous tubules (Shima et al., 2013). Studies have shown that Leydig cell differentiation undergoes immature-mature stage (Zhou et al., 2019). We also identified immature and mature Leydig cells. Re-clustering of Leydig cells generated twelve distinct cell clusters, including ten iLeydig cells and two Leydig cells (Figure S9A). Heatmap showed specific expression of markers in Leydig cells (Figure S9B). RSPO3 and MYOCD were highly expressed in iLeydig cells. CYP17A1 and CCDC69 were highly expressed in Leydig cells. Leydig cells in different samples are shown in Figure S9C. OA samples had more iLeydig cells than NOA samples (Figure S9D).

### Peritubular myoid cells (PMCs) heterogeneity and development

PMCs are the key cellular components of the wall of seminiferous tubules and they are thought to be of great importance for the intratesticular transport of immotile sperm (Wang, Chen, & Liu, 2018). Re-clustering of PMCs revealed that there were 10 PMCs subclusters (Figure S10A). Heatmap showed specific expression of markers in PMCs (Figure S10B). PMCs cells in different samples are shown in Figure S10C. NOA samples had more PMCs-1 than OA samples (Figure S10D). PMCs-1 might play an important role in spermatogenesis.

### Macrophage heterogeneity and development

Macrophages play an important role in spermatogenesis and can produce 25-hydroxycholesterol within the testosterone biosynthetic pathway, thus contributing to the testosterone production process (Potter & DeFalco, 2017). Re-clustering of macrophage revealed that there were five macrophage subclusters (Figure S11A). Heatmap showed specific expression of markers in macrophage (Figure S11B). Macrophage cells in different samples are shown in Figure S11C. NOA samples had more macrophage-2 than OA samples (Figure S11D). Macrophage-2 might play a key part in spermatogenesis.

## Discussion

NOA is due to pretesticular factors or testicular factors, usually abnormal spermatogenesis, which cannot be cured by surgery (Zarezadeh et al., 2021). Therefore, it is particularly important to study the etiology of NOA. Spermatogenesis is a complex developmental process that requires coordinated differentiation of multiple cell lines. In present study, we performed scRNA-seq and scATAC-seq to assess the heterogeneity of germ cells and somatic cells. We identified 12 subtypes for germ cells, 8 subtypes for Sertoli cells, 12 subtypes for Leydig cells, 10 subtypes for PMCs, 8 subtypes for macrophage and marker genes of specific cell type. The process of spermatogenesis was determined based on the cell trajectory analysis. Collectively, our results provide rich resources for exploring the potential mechanism of spermiogenesis.

In this study, some specific marker genes for germ cells were screened. We found that UTF1 and ID4 were specifically expressed in SSC1, and DMRT1 was specifically expressed in Diffing SPG. ID4 was a key regulator of SSC and ID4 also marked SSCs in the mouse testis (Sun, Xu, Zhao, & Degui Chen, 2015). UTF1 was reported to be a marker for SSCs in stallions (Jung, Roser, & Yoon, 2014). UTF1 was discovered to play an important part in germ cell development, spermatogenesis, and male fertility in mice (Raina, Dey, Thool, Sudhagar, & Thummer, 2021). DMRT1 is necessary for male sexual development (Zhang, Oatley, Bardwell, & Zarkower, 2016). The mechanism and expression pattern of these marker genes still need further research.

SSC are adult stem cells in the testis of mammals that maintain spermatogenesis and are essential for male fertility, but the mechanisms remain elusive. UTF1 is a transcription factor expressed in SSC1, which plays a definite role in the proliferation and differentiation of pluripotent stem cells (Tan & Wilkinson, 2019). Dmrt1 belongs to a family of conserved transcriptional regulators that control several key processes in mammalian testis, including germ cells and somatic cells. A recent discovery suggested that DMRT1 was essential for the formation of SSC and had gonadal-specific and amphoteric dimorphic expression patterns (Sohni et al., 2019). NANOS3 is a member of the highly conservative NANOS family. The decrease in NANOS3 expression could result in a decrease in the number of germ cells (Julaton & Reijo Pera, 2011). In this study, UTF1 was specifically expressed in SSC1, and DMRT1 and NANOS3 were the main specifically expressed genes in SSC2. Therefore, UTF1, DMRT1 and NANOS3 might promote SSC proliferation and was necessary for SSC differentiation.

Spermatogenesis consists of three processes: mitosis of spermatogonia, meiosis of spermatocyte and deformation of spermatocyte. Abnormalities in any of these stages can result in spermatogenesis disorders. During this process, the testicular microenvironment in which spermatogenic cells are located plays an important role. In addition to Sertoli cells, the testicular microenvironment also includes myoid cells and mesenchymal cells and their secreted factors (Zhou et al., 2019). In this study, we clustered four somatic cell types: Sertoli cells, Leydig cells, PMCs and macrophages. Sertoli cells play an important role in normal spermatogenesis (Zomer & Reddi, 2020). They provide the physical framework that supports the survival of germ cells and secrete unique growth factors and cytokines to assist in the development of germ cells (O’Donnell, Smith, & Rebourcet, 2022; Wu, Yan, Ge, & Cheng, 2020). In our study, the NOA3 group had the least number of sperm cells due to the lack of SC3/8, and the NOA2 group had a smaller number of SC3 cells than the normal OA group, so the number of sperm cells in the NOA2 group was between the NOA3 and OA groups. Although NOA2 did not have SC8 to participate in the formation of SPT3, this process was compensated by SC3, which was involved in cell formation throughout all stages of sperm development, and its function was particularly important. These data suggested that the association between germ cells and Sertoli cells was stage-specific, but this “specificity” is not entirely isolated.

Notch receptors have been reported to play an important role in cell fate determination, maintenance and differentiation of stem cells (Huang, Rivas, & Agoulnik, 2013). Notch signaling components have been discovered to express in Sertoli and germ cells in the developing and mature testis (Garcia, DeFalco, Capel, & Hofmann, 2013). In mice, a breakdown of Notch signaling can lead to abnormal cell differentiation and early embryonic lethality (McCright et al., 2001; Swiatek, Lindsell, del Amo, Weinmaster, & Gridley, 1994). Murta et al. (2014) also found notch signaling disrupt results in abnormal spermatogenesis in the mouse. Notch signaling was reported to be a key pathway regulating Sertoli cell physiology, and its changes may disturb reaction of Sertoli cells to androgens (Kaminska et al., 2020). Notch signaling in Sertoli cells is indirectly associated with germ cell development (Lu et al., 2019). The upregulation of notch signaling in Sertoli cells induced the transformation the quiescence to differentiation and meiosis in germ cells (Garcia & Hofmann, 2013). In this study, we found Notch signaling were involved in the interaction between germ cells and Sertoli cells. Therefore, Notch signaling might function in spermatogenesis.

In conclusion, our results revealed 12 germ cell subtypes and 8 Sertoli cell subtypes. We determined the process of spermatogenesis and found that SC3 subtypes (marked by SPATS1) of Sertoli cell played an important role in this process. The interaction between germ cells and sertoli cells at each differentiation stage were stage-specific. Notch1/2/3 signaling were discovered to be involved in germ cell-Sertoli cell interaction. Our results not only give us a comprehensive insight into human spermatogenesis, but also pave the way for determining molecules participated in the development of male germ cells, offering a powerful tool for further study on NOA.

## Acknowledgements

Not applicable.

## Author contributions

SW: Conceptualization, Data curation, Writing—original draft; HW: Conceptualization, Data curation, Writing—original draft; BJ: Visualization, Formal analysis; QZ: Visualization, Formal analysis; HY: Methodology, Project administration; DZ: Supervision, Project administration, Writing—review & editing. All authors read and approved the final manuscript.

## Data Availability Statement

ScRNA-seq data have been deposited in the NCBI Gene Expression Omnibus with the accession number GSE202647, and scATAC-seq data have been deposited in the NCBI database with the accession number PRJNA1177103.

## Ethics Approval and Consent to Participate

This study was approved by the Ethics Committee of Renji Hospital, Shanghai Jiao Tong University School of Medicine. All patients provided written informed consent.

## Supplementary Information

**Figure S1. scATAC-seq analysis of the entire cell subpopulation.** (A) UMAP analysis of 10 cell subsets in scATAC-seq results. (B) Accessibility peak plots of markers for cell subpopulations.

**Figure S2. Integrating scRNA-seq and scATAC-seq analysis.** (A-C) Integrating all cell types (A), germ cell types (B), Sertoli cell types (C) from scRNA-seq and scATAC-seq by UMAP plot.

**Figure S3. Analysis of germ cell markers in spermatogenesis.** (**A**) Heatmap showed marker gene expression of germ cell subtypes. (**B**) Gene expression patterns of germ cell markers on violin plots. (**C**) Distribution diagram of germ cell subsets using pseudotime analysis.

**Figure S4. scATAC-Seq Clustering.** (A) Heatmap of germ cell markers in scATAC-seq results. (B) Heatmap of motifs in scATAC-seq results.

**Figure S5**. **Germ cell function analysis and epigenetic analysis.** (**A-C**) Analysis of beam cluster transcription factors and its GO term. (**D**) Heatmap of transcription factors in germ cells. (**E**) GO analysis of different states of Sertoli cells. (**F**) Qusage analysis of different states of Sertoli cells.

**Figure S6. GO analysis of germ cells.**

**Figure S7. GO analysis of Sertoli cells.**

**Figure S8. Correlation analysis of germ and Sertoli cell expression.** (**A**) The interaction between germ cells and Sertoli cells through Notch signaling pathway. (**B**) Accessibility peak plots of NOTCH1/2/3.

**Figure S9. Identification of Leydig cell types in the testis.** (**A**) Clustering on the Leydig cells revealed 12 different cell types. (**B**) Heatmap showed Leydig cell subtype specific maker genes. (**C**) The distribution of Leydig cells in different samples. (**D**) Proportion of Leydig cell subtypes in each sample.

**Figure S10. Identification of PMCs types in the testis.** (**A**) Clustering on the PMCs revealed 8 different cell types. (**B**) Heatmap showed PMCs subtype specific maker genes. (**C**) The distribution of PMCs in different samples. (**D**) Proportion of PMCs subtypes in each sample.

**Figure S11. Identification of macrophage types in the testis.** (**A**) Clustering on the macrophage revealed 6 different cell types. (**B**) Heatmap showed macrophage subtype specific maker genes. (**C**) The distribution of macrophage in different samples. (**D**) Proportion of macrophage subtypes in each sample.

**Table S1 The number of different kinds of cells in five samples in scRNA-seq. Table S2 The number of different kinds of cells in five samples in scATAC-seq.**

