## Supplementary figures and images for "ScRNA-seq and scATAC-seq reveal that sertoli cell mediate spermatogenesis disorders through stage-specific communications in non-obstructive azoospermia"

### Figure S1

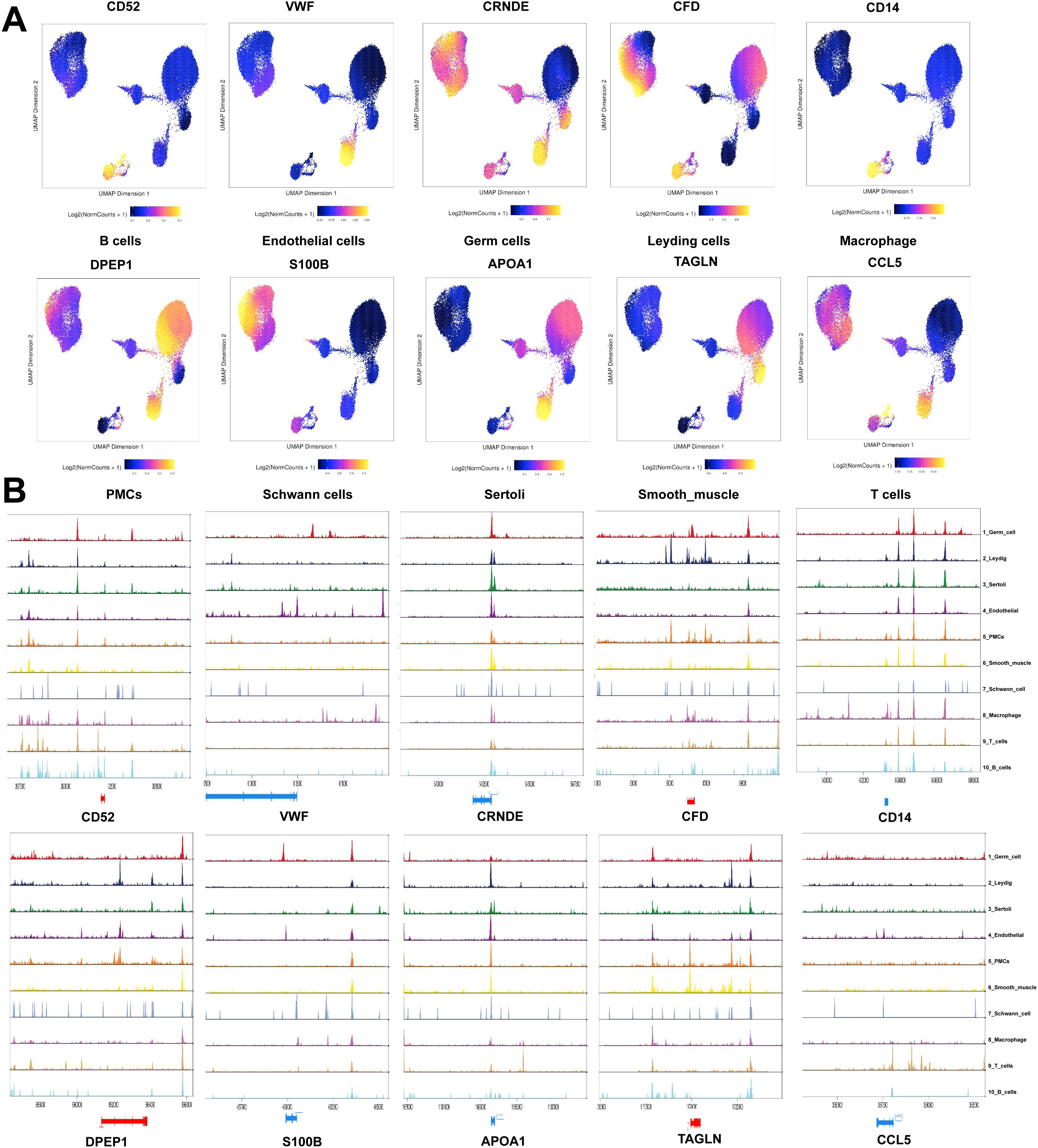

### Figure S2

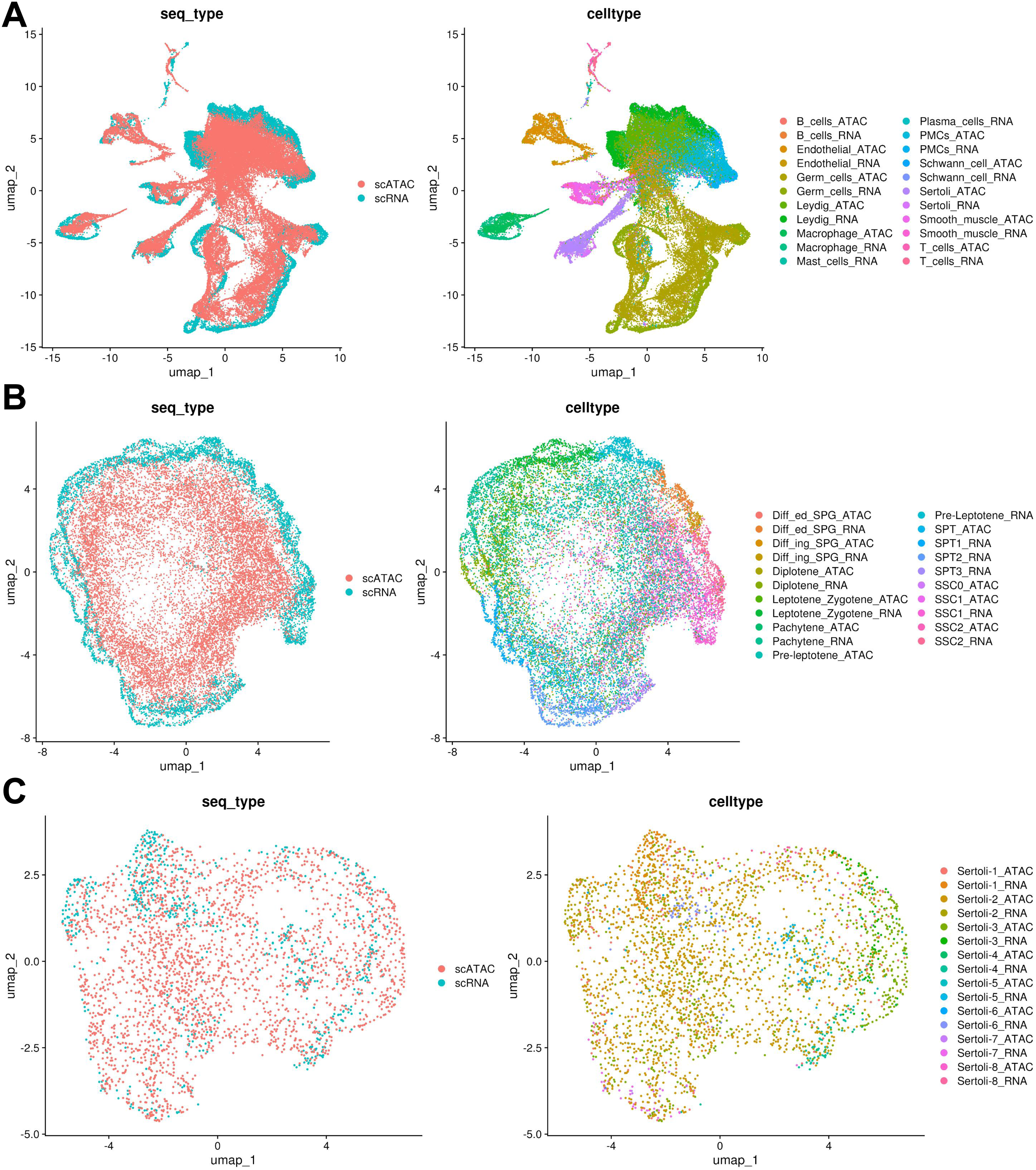

### Figure S3

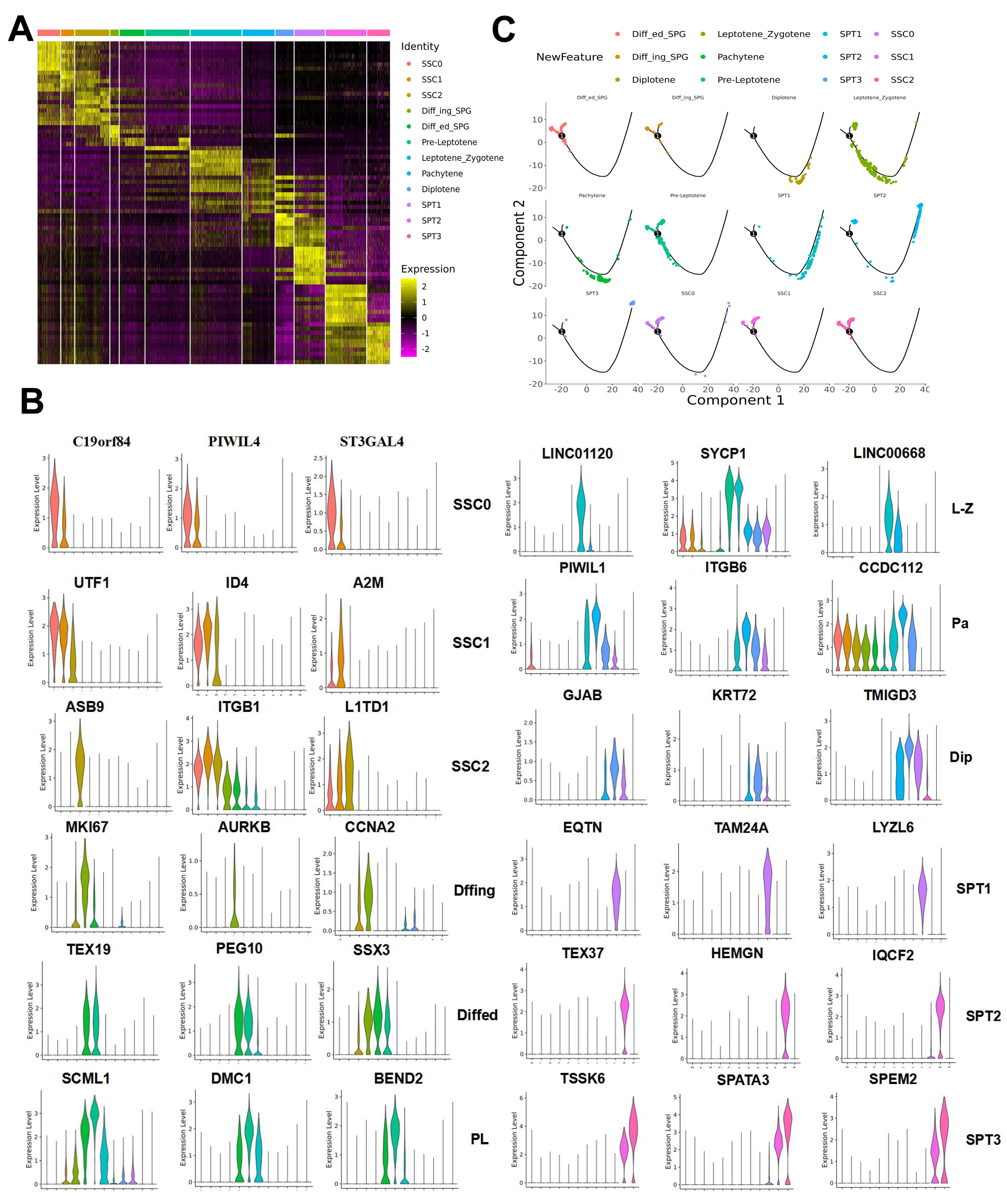

### Figure S4

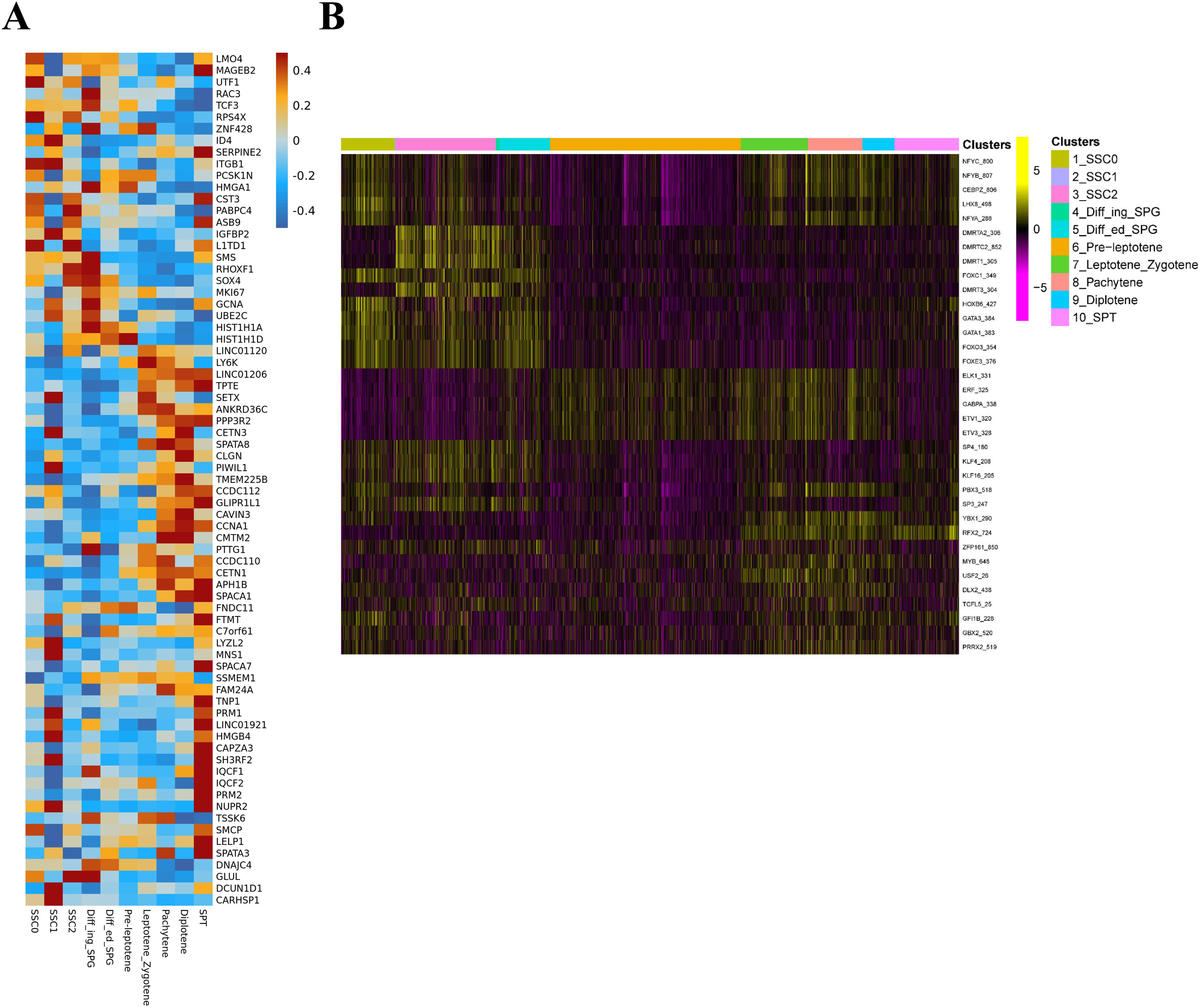

### Figure S5

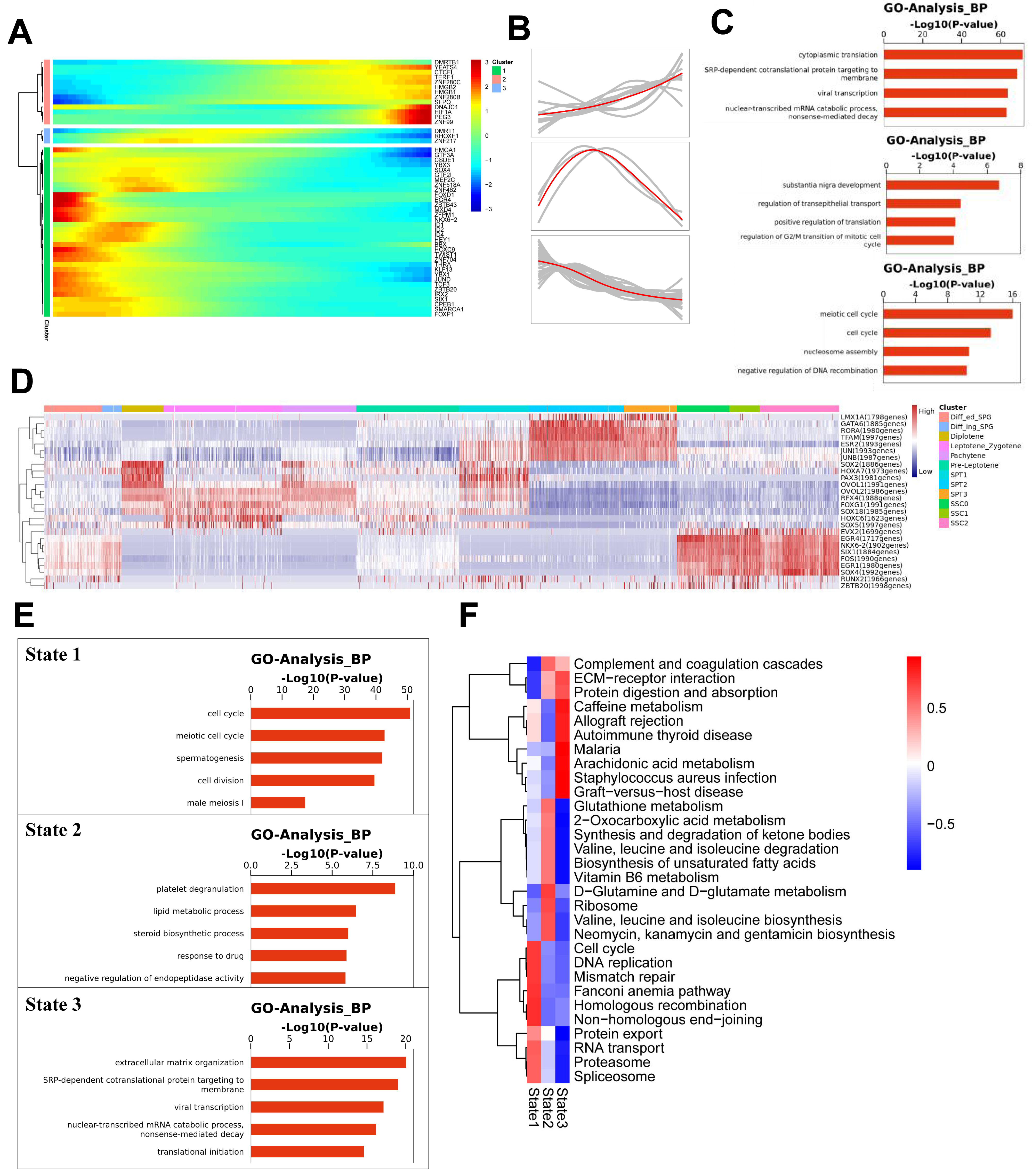

### Figure S6

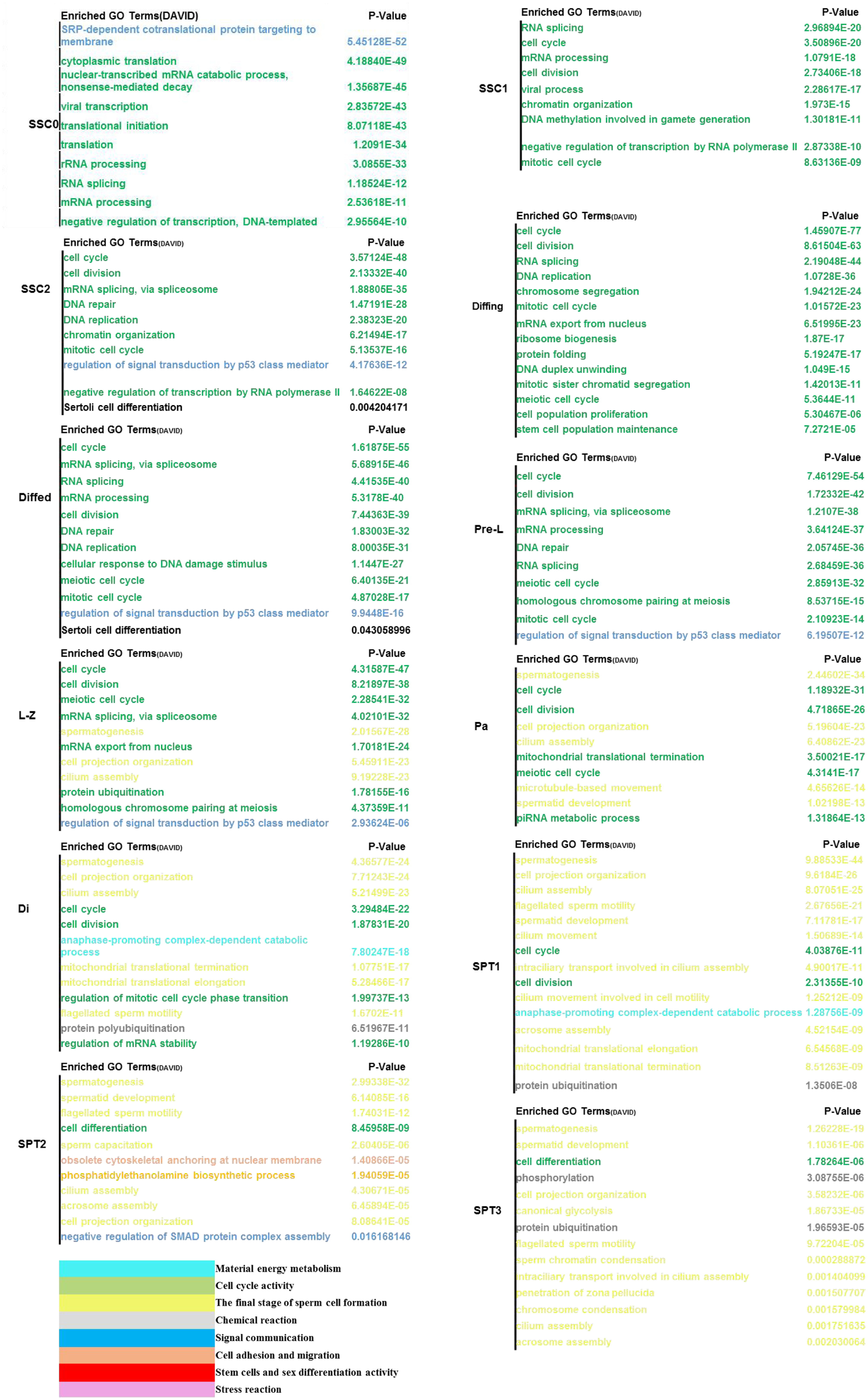

### Figure S7

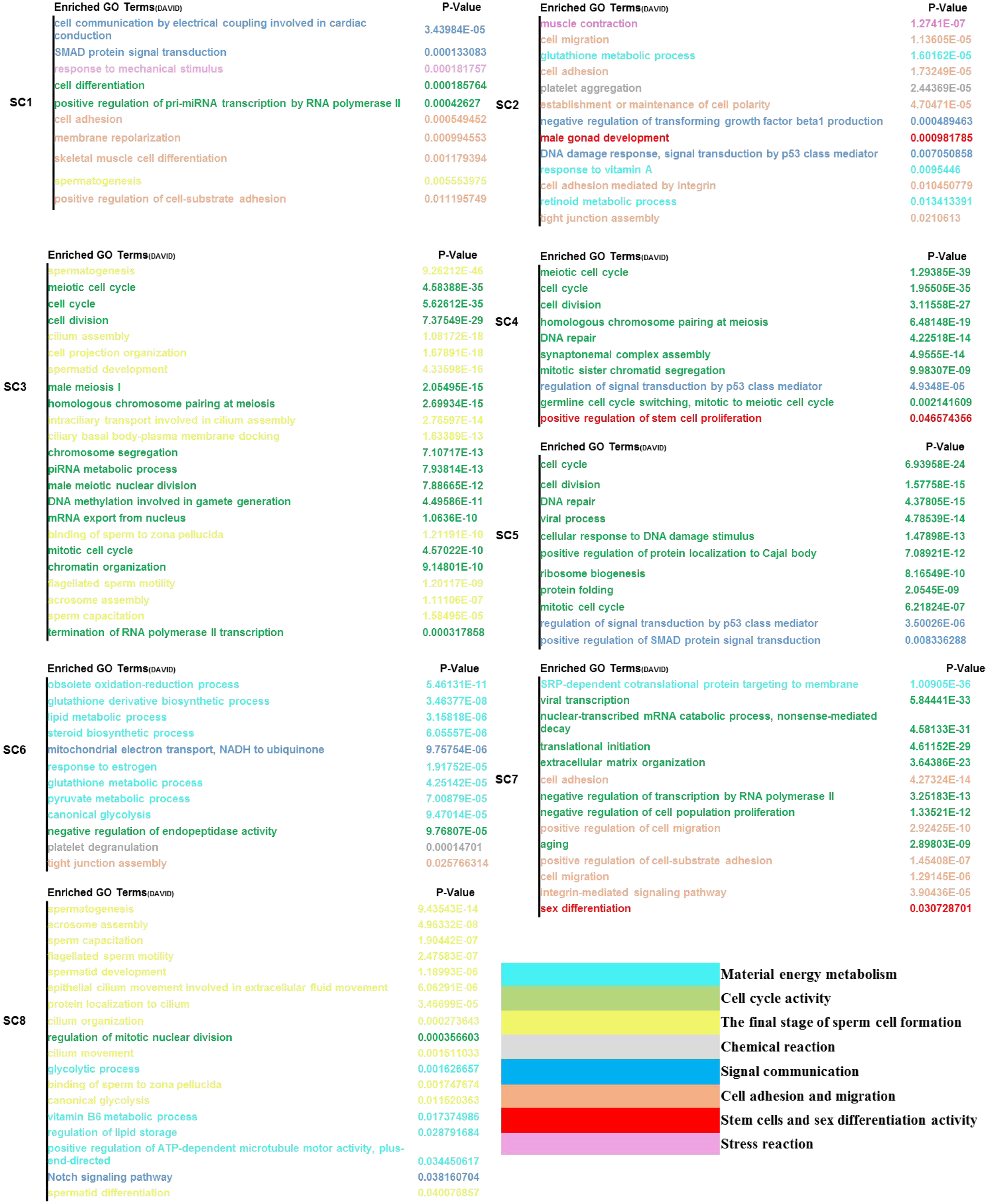

### Figure S8

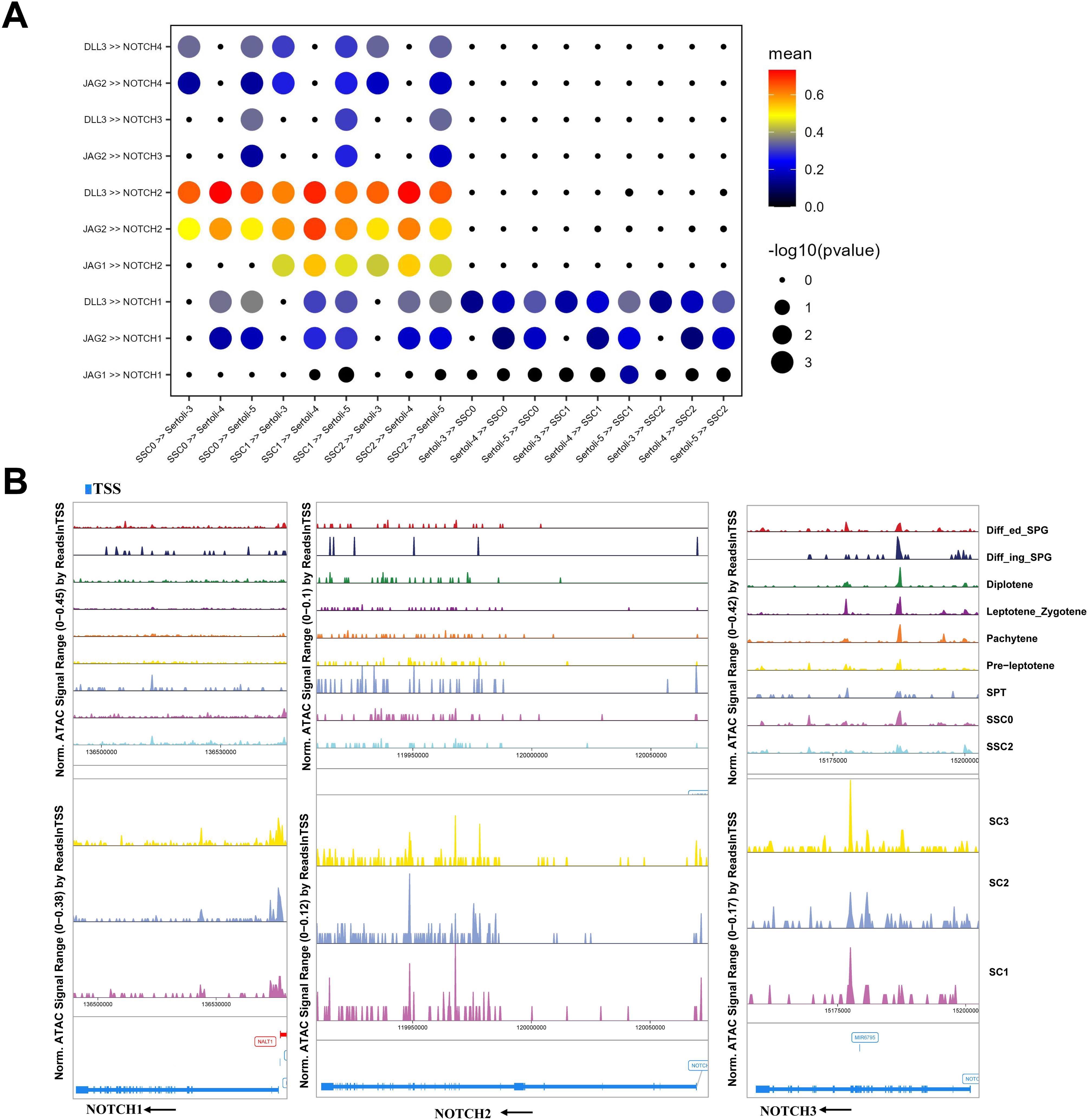

### Figure S9

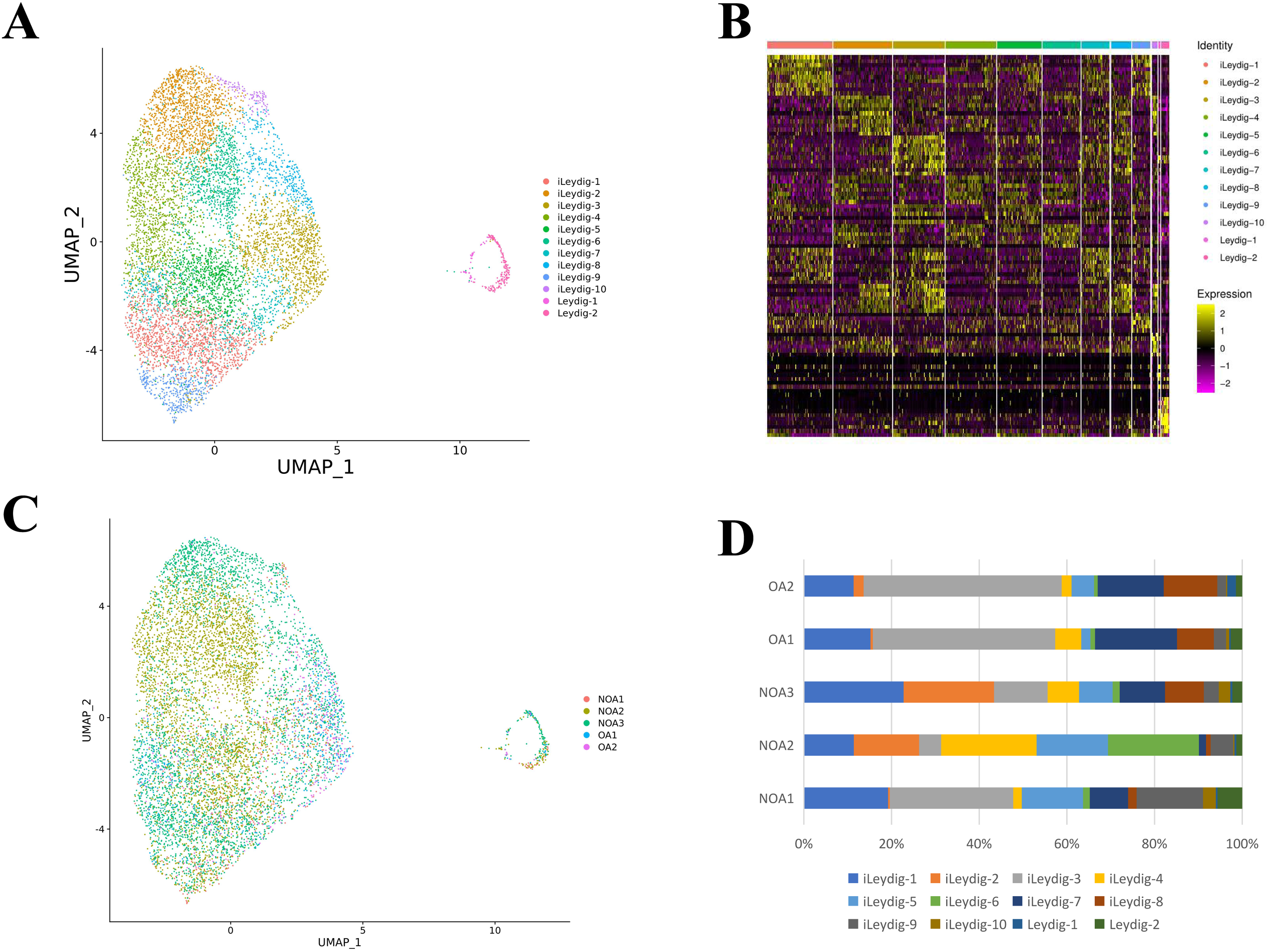

### Figure S10

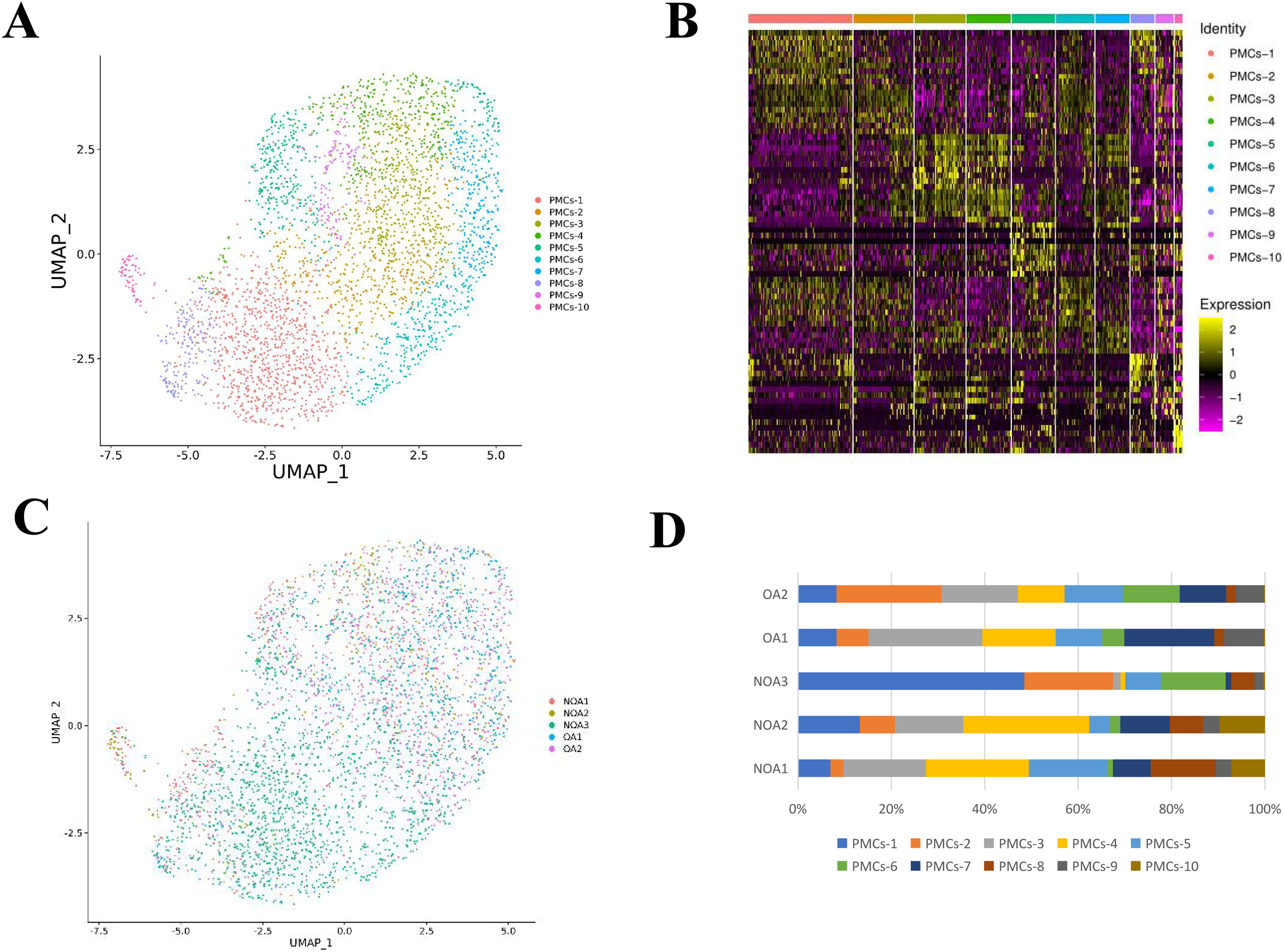

### Figure S11

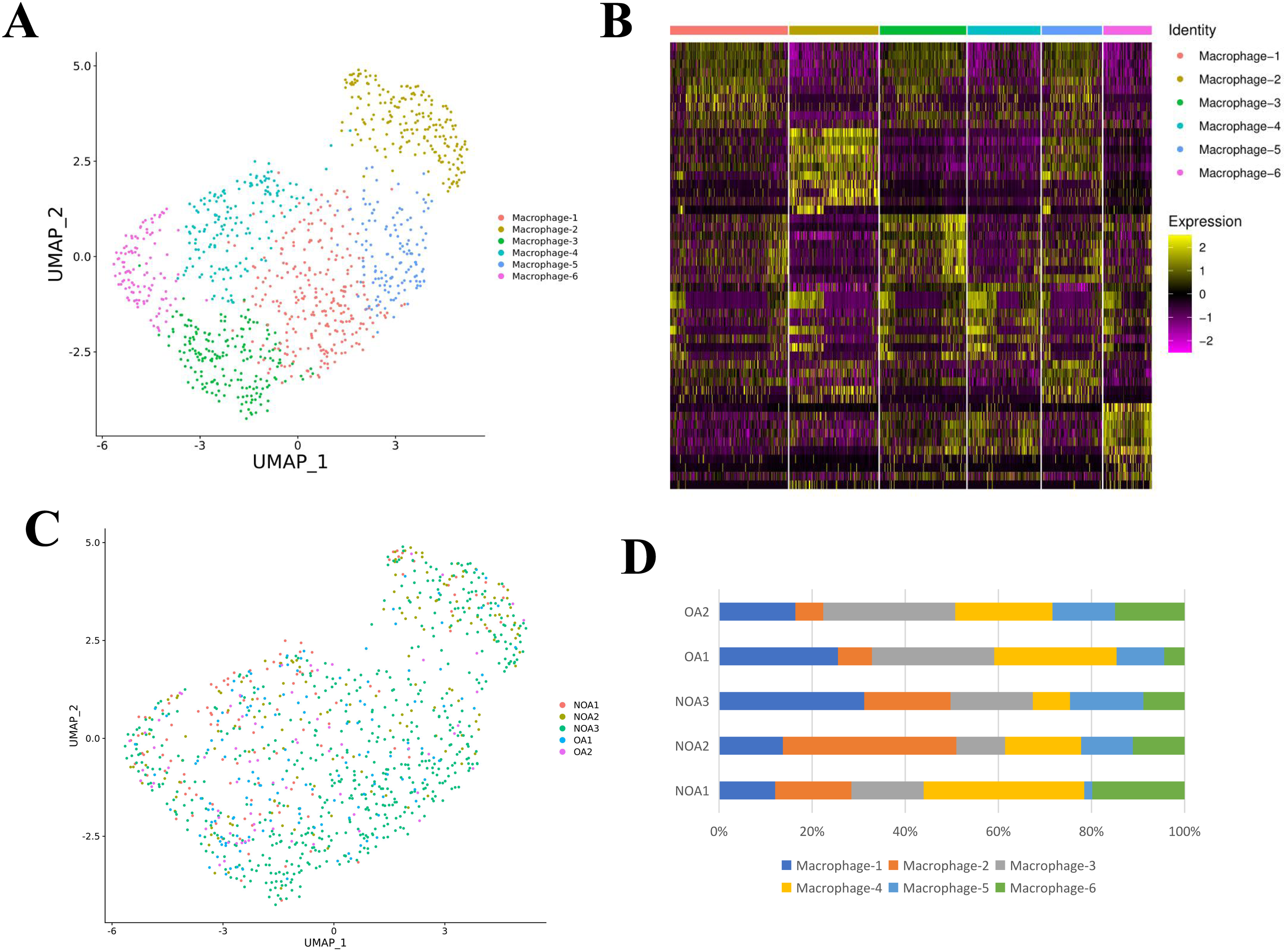
