## Supplementary material for "ScRNA-seq and scATAC-seq reveal that sertoli cell mediate spermatogenesis disorders through stage-specific communications in non-obstructive azoospermia": Table S1

| **Cluster** | **NOA1** | **NOA2** | **NOA3** | **OA1** | **OA2** |
| --- | --- | --- | --- | --- | --- |
| Germ_cells | 1654 | 792 | 0 | 2329 | 2124 |
| Leydig | 515 | 3892 | 3625 | 572 | 352 |
| Sertoli | 90 | 22 | 123 | 239 | 286 |
| Endothelial | 108 | 223 | 188 | 150 | 224 |
| PMCs | 544 | 226 | 1427 | 701 | 641 |
| Smooth_muscle | 278 | 380 | 325 | 108 | 212 |
| Schwann_cell | 6 | 22 | 37 | 3 | 11 |
| Macrophage | 116 | 153 | 577 | 137 | 67 |
| Mast_cells | 2 | 11 | 17 | 6 | 0 |
| T_cells | 19 | 110 | 153 | 15 | 15 |
| B_cells | 0 | 22 | 4 | 0 | 1 |
| Plasma_cells | 0 | 33 | 2 | 0 | 0 |

**Table S1** **The number of different kinds of cells in five samples in scRNA-seq.**
