## Supplementary material for "ScRNA-seq and scATAC-seq reveal that sertoli cell mediate spermatogenesis disorders through stage-specific communications in non-obstructive azoospermia": Table S2

**Table S2** **The number of different kinds of cells in five samples in scATAC-seq.**

| **Cluster** | **NOA1** | **NOA2** | **NOA3** | **OA1** | **OA2** |
| --- | --- | --- | --- | --- | --- |
| Germ_cell | 4734 | 630 | 0 | 4548 | 3823 |
| Leydig | 1757 | 4662 | 4271 | 902 | 625 |
| Sertoli | 474 | 32 | 280 | 1022 | 827 |
| Endothelial | 848 | 987 | 443 | 387 | 565 |
| PMCs | 1665 | 405 | 1419 | 862 | 1024 |
| Smooth_muscle | 968 | 577 | 266 | 109 | 276 |
| Schwann_cell | 37 | 17 | 9 | 2 | 5 |
| Macrophage | 451 | 273 | 722 | 253 | 126 |
| T_cells | 59 | 112 | 147 | 19 | 18 |
| B_cells | 2 | 60 | 7 | 0 | 1 |
